# Tumoral switch in NUMB splicing changes essential transcription pathways and induces malignant properties in tumour cells

**DOI:** 10.64898/2026.05.15.725391

**Authors:** José Manuel García-Heredia, Sara M. Ortega-Campos, Amancio Carnero

**Affiliations:** Instituto de Biomedicina de Sevilla (IBIS)/HUVR/CSIC/Univ Sevilla, Hospital Universitario Virgen del Rocío, Ed. IBIS, Avda. Manuel Siurot S/N, 41013, Seville, Spain; CIBER de Cancer (CIBERONC), Instituto de Salud Carlos III, Madrid, Spain; Departamento de Bioquímica Vegetal y Biología Molecular, Universidad de Sevilla, 41012 Sevilla, Spain

**Author notes:** Corresponding authors, Instituto de Biomedicina de Sevilla (IBIS)/HUVR/CSIC/Univ Sevilla, Hospital Universitario Virgen del Rocío, Ed. IBIS, Avda. Manuel Siurot S/N, 41013, Seville, Spain. These authors contributed equally (José Manuel García-Heredia, Sara M. Ortega-Campos).

**Keywords:** NUMB, breast cancer, transcript isoform balance, transcriptomics, pharmacogenomics, TCGA, CCLE, patient-derived xenografts

## Abstract

**Background:** Recent evidence suggests that cancer can exhibit splicing alterations that give rise to tumour-specific isoforms. One example is NUMB, which produces four isoforms (p72, p71, p66, and p65) through alternative splicing of exons 3 and 9. Traditionally considered a tumour suppressor, it also has been considered an oncogene. We propose that this duality is due to isoform-specific expression.

**Results:** Using public databases, we identified a tumour-associated switch in NUMB isoform expression: p72/p71 are upregulated in tumours, whereas p66/p65 are more expressed in non-tumour tissues. These isoforms correlate differently with cellular processes. NUMBL, a NUMB homolog, behaves similarly to p65. We identified two transcriptional clusters: one characterized by high expression of p72/p71, and another by p66/p65/NUMBL. Each group was associated differently with the Notch, WNT/β-catenin, Hedgehog, and Hippo signalling pathways, suggesting isoform-specific regulatory roles. Analysis of breast cancer cell lines (CCLE) led to a NUMB score based on isoform expression, which classified cell lines into biologically distinct groups. The p72/p71-enriched group showed distinct signatures, pathway activity, and drug sensitivity. Applying this score to TCGA-BRCA samples revealed a significant link between high NUMB-score and poor survival, confirmed by Kaplan–Meier analysis.

**Conclusions:** NUMB emerges as a potential oncogenic contributor and biomarker in splicing-based personalised medicine, highlighting isoform-specific expression as a clinically relevant determinant of tumour behaviour, pathway activity, and therapeutic response.

## INTRODUCTION

The increase in knowledge based on genetic diagnosis is allowing several branches of medicine to adopt more personalized strategies, leaving behind approaches based on the symptoms. These strategies combine this information with clinical data, obtaining more targeted therapies, with increased efficacy, that allow a better prediction of disease progression. Among the multiple factors underlying why some patients with a specific disease respond to the treatment, while for other patients the same treatment has no effects, a molecular mechanism gaining visibility, for its key contribution, is Alternative Splicing (AS). This evolutionarily conserved mechanism in multicellular organisms boosts molecular complexity by allowing a single gene to give rise to multiple mRNA isoforms and expand the variety of transcripts and proteins available. Splicing is a process that affects mRNA maturation, carried out by a complex machinery composed of small nuclear RNAs and more than 300 proteins. While one of its main functions is intron removal and the subsequent exon junction, it also influences other stages of gene expression, such as mRNA export, translation, and the degradation of defective transcripts through quality control mechanisms. These regulations are essential for maintaining cellular homeostasis. It is estimated that more than 90% of human genes undergo AS, and many resulting isoforms encode proteins that differ in their interaction partners and functional behaviour. Although often assumed to be similar, splice variants can have distinct effects on cellular processes. The selection of specific isoforms is tightly regulated and varies across cell types and developmental stages. As a result, certain biological programs, such as neural development, epithelial-to-mesenchymal transition (EMT), and T-cell activation exhibits different versions of a gene, due to changes in AS. These differences in AS patterns between cell types represent a valuable source of diagnostic and therapeutic biomarkers [1–8].

Studying how different isoforms of a gene diverge functionally can help us understand certain factors that control gene expression [9], such as RNA regulatory elements and the activity of transcription factors. Different strategies to modulate splicing include, for example, the inhibition of a specific spliceosomal component or the inhibition of a regulatory splicing factor. However, because many elements of the splicing machinery are shared between healthy and malignant cells, such interventions must be approached with caution and thoroughly evaluated to avoid unintended toxicity. Knowing how these processes work would also provide information on the molecular basis of different diseases caused by aberrant splicing and thus propose possible more effective therapies against them [10]. The essential role of mRNA splicing as a general process means that alterations to the splicing machinery, such as missense mutations in splicing factors or changes in their expression levels, can be deleterious or even lethal. Such aberrations can lead to organ dysfunction, particularly when they affect tissue-specific genes and alter cell population dynamics. Aberrant AS has been connected to a broad range of diseases, such as cancer, heart disease, diabetes, Alzheimer’s, Huntington’s, celiac disease, psoriasis and lupus, among others [11, 12]. Among its consequences are intron retention, inappropriate activation of early-development isoforms, and mutations that disrupt normal splice sites, sometimes leading to the inactivation of tumour suppressors or the emergence of oncogenic forms. These events often result from mutations in regulatory sequences or disruptions in the activity of splicing factors. Among the most frequently affected components of the spliceosome are U2AF1, SRSF2, and SF3B1, appearing the latter mutated in a significant percentage of myelodysplastic syndromes [13].

In cancer, there is increasing evidence that AS dysregulation could be included into the set of traits increasingly considered as hallmarks of cancer [14–16]. This dysregulation would lead to a change in the isoform expression patterns of certain genes, favouring the production of isoforms that promote proliferation, inhibit cell death, stimulate angiogenesis, enhance invasion and metastatic dissemination, generate drug resistance or reprogram cell metabolism [3, 14–17]. The presence of specific AS events in tumours highlights the need for increased research into the mechanisms that lead to such aberrant AS, as well as the discovery of potential therapeutic applications [18, 19]. Changes in isoform expression that occur not only during oncogenic transformation but also in other diseases could serve as biomarkers to stratify patients who might benefit from therapies targeting splicing regulation [19–21]. These strategies could include inhibiting splicing factors with chemotherapy, regulating certain splicing events with oligonucleotides [22], developing immunotherapies based on neoantigens derived from cancer-specific AS events, or identifying oncogenes with splicing-associated therapeutic vulnerability [19, 23–25]. Indeed, some of these cancer-specific splicing events have also been linked to increased resistance to targeted therapies, reinforcing their potential as clinically relevant biomarkers.

One potential isoform-specific biomarker is NUMB. Originally identified in *Drosophila melanogaster*, *NUMB* is a highly conserved gene with a close vertebrate homologue, *NUMBL* [26, 27]. *NUMB* consists of ten exons and produces up to nine different transcripts through AS. Among these, only four are protein-coding isoforms, generated by the inclusion or exclusion of exons 3 and 9. These isoforms are named based on their molecular weight: variant 1 (p72), variant 2 (p66), variant 3 (p71) and variant 4 (p65) [28–30]. Interestingly, NUMB participates in multiple signalling pathways by acting as a charge-selective adaptor [31–34]. In the NOTCH pathway, recruitment of the E3 ubiquitin ligase ITCH by NUMB promotes ubiquitination of the Notch intracellular domain (NICD), resulting in its degradation in the proteasome. This prevents NICD from being transported to the nucleus, thus reducing the activity of the NOTCH pathway [32, 35, 36]. Moreover, NUMB is also involved in the Hedgehog (SHH) pathway, essential in both development and tumorigenesis [37]. NUMB acts as a suppressor of this pathway by recruiting the E3 ubiquitin ligase ITCH, which induces the degradation of GLI1 in the proteasome, thus preventing pathway activation [38]. In addition, NUMB also appears to play a role in the WNT/β-catenin pathway and may itself be a downstream target of its canonical pathway, so that when this pathway is activated, it triggers the β-catenin cascade, inducing the expression of NUMB [31, 38]. There also appears to be a regulatory feedback loop between both proteins, as NUMB has been shown to promote proteasomal degradation of β-catenin, thus preventing pathway activation [39–41]. However, and although some functions have been ascribed more to a certain isoform, much of the published results take NUMB, and NUMBL, as a unique “protein”, with their isoforms sharing functions, which could be an oversimplification of what happens in the cell.

In the present work, NUMB was studied as a potential tumorigenic biomarker in the context of personalised medicine strategies targeting splicing events. Likewise, we showed the differential regulation that these isoforms could exert on different cellular pathways involved in tumorigenesis, as well as the *in vivo* phenotypic changes that could be associated with the different expression of the isoforms. Based on our analyses, we propose that the in-depth study of NUMB isoforms could eventually allow the creation of patient profiles for the development of personalised cancer therapies. The overall analytical workflow of the study is shown in **Supplementary Figure S1**.

## RESULTS

### The splicing process undergoes an oncogenic switch in tumorigenesis

To explore the role of AS in cancer, we analysed isoform switch events across multiple TCGA tumour types compared to non-tumour tissue. Splicing alterations were observed in all tumour types analysed (**Figure 1A**), finding that the most frequent types of splicing change across all tumours were Alternative Termination site (AT) and Exon Skipping (ES) (**Figure 1B**). To further analysis, we chose to focus on breast tumours, as previous studies had reported reduced NUMB expression in these tumours [42–44]. We performed a detailed analysis of the different types of splicing changes between non-tumour and tumour tissue, specifically in breast cancer, finding increments in the number of events in tumour samples for Alternative 3’ (A3), Alternative Transcription Start Site (ATSS), Alternative Transcription Termination Site (ATTS) and Exon Skipping (ES). On the other hand, a decrease in the number of Mutually Exclusive Exons (MES), Intron Retention (IR) and Alternative 5’ (5A) events was observed compared to non-tumour tissue (**Figure 1C**). These data indicate marked differences in alternative splicing patterns between tumour and non-tumour breast tissue, which may contribute to oncogenic transformation.

**Figure 1.**
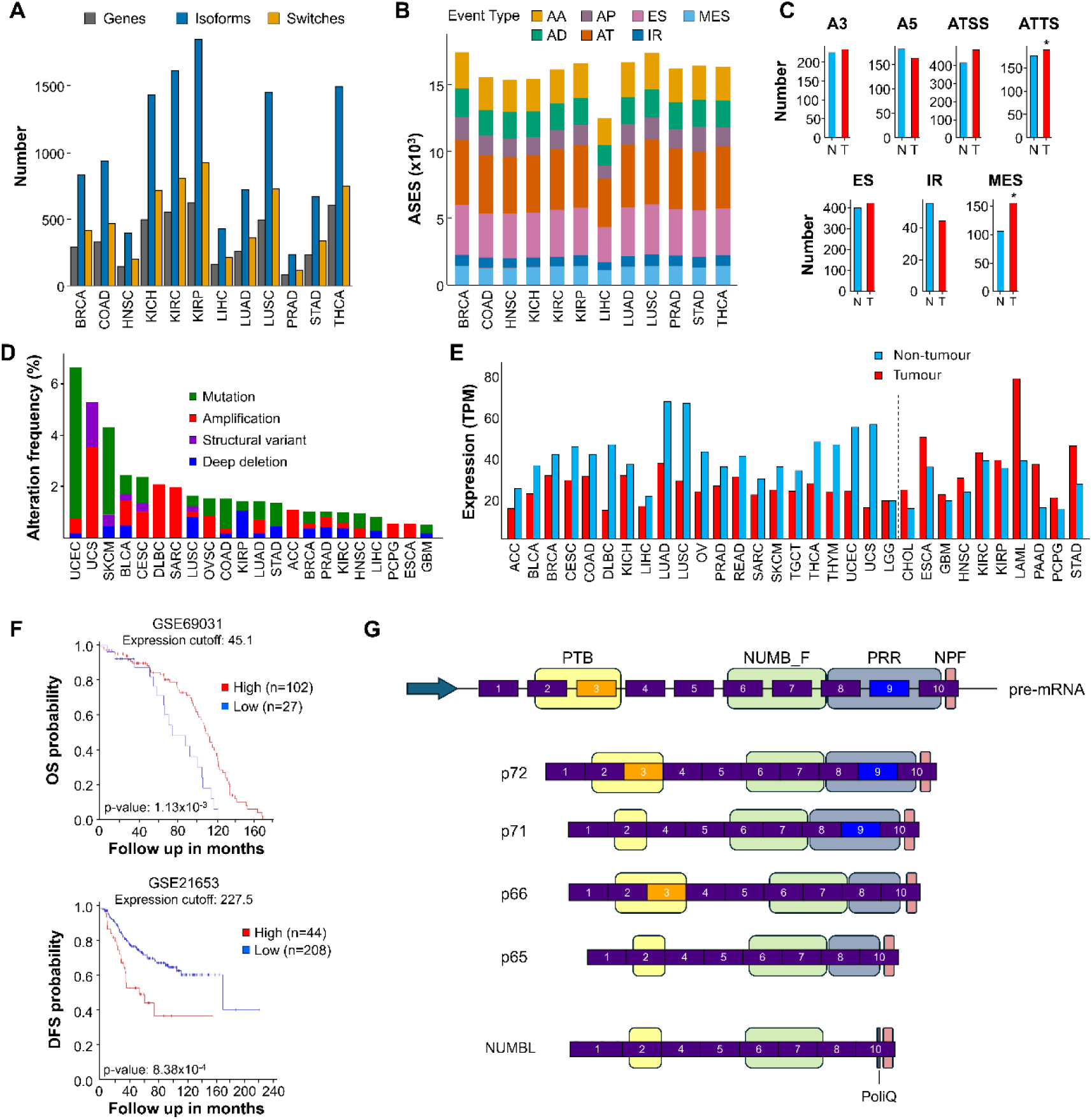
The splicing process undergoes an oncogenic switch. **A)** Study of the number of isoforms, switches, and genes with AS alterations across different tumour types from the TCGA database. **B)** Distribution of AS event (ASE) types observed in the different TCGA tumours. **C)** Comparison of splicing events observed between non-tumour and tumour tissue in breast cancer. Directional bias (gain vs loss) was assessed using binomial tests with Benjamini–Hochberg correction. ATSS (FDR = 0.044) and MES (FDR = 0.016) showed significant directional bias (*FDR < 0.05). **D)** Frequency and type of NUMB alterations in cancer, including mutations, structural variants, amplifications and deep deletions, based on data from the TCGA database. Tumours are ordered from highest to lowest alteration frequency. **E)** Differential expression of NUMB in non-tumour tissue compared to tumour. Dashed line marks the separation between tumours with higher NUMB expression compared to normal tissue and those with lower expression. Splicing event classes and TCGA tumour abbreviations are provided in **Supplementary Table S1**. **F)** Kaplan-Meier plot showing the association between NUMB expression and survival probability in breast datasets GSE69031 and GSE21653. OS: Overall survival; DFS: Disease Free Survival. **G)** NUMB generates four isoforms through AS of exons 3 and 9. The absence of these exons modifies the PTB (phosphotyrosine domain) and PRR (proline rich region) domains of the protein. Other domains, such as the NUMB-F domain and the NPF (Asn-Pro-Phe) motif remain unchanged. NUMBL shares high structural similarity with isoform p65, except for the presence of a PolyQ (polyglutamine) domain in NUMBL. Exon numbering exons is based on the NUMB gene.

cBioportal was used to analyse the possible presence of alterations in the NUMB gene that could contribute to tumorigenesis [45]. Several tumour types exhibited low to moderate frequencies of NUMB alterations, with uterine endometrial carcinoma showing the highest rate, around 6.5%. The most common alterations include mutations in the gene sequence or amplifications, increasing the gene copy number. Less frequently, structural variants or deep deletions were also detected (**Figure 1D**). The low percentage of alterations detected in BRCA tumours, around 1%, suggested that the effects we observed were more likely due to modifications in expression patterns, such as AS, rather than to the expression of a mutated version of NUMB.

Subsequently, we compared NUMB expression in healthy tissues versus different tumours using the GEPIA2 database [46]. The results revealed marked differences in NUMB expression between non-tumour and tumour tissue across all tissue types analysed (**Figure 1E**). Of the 31 tumour types analysed, 10 showed higher NUMB expression in tumours compared to non-tumour tissue (approximately 32%). In the remaining tumour types, NUMB expression was comparable to or lower than in non-tumour tissue, consistent with a context-dependent role of NUMB across cancers. Similarly, analysis of patient data revealed significant variability in survival outcomes. In some cases, high NUMB expression correlated with improved survival, while in others it was associated with poorer prognosis (**Figure 1F, Supplementary Figure 2**).

The differences observed both in NUMB expression levels across tumour types and in survival outcomes, as shown by the Kaplan–Meier analysis, do not allow us to clearly define NUMB as either oncogenic or tumour suppressive. This dual behaviour could be due to multiple causes, including the tissue of origin, differential methylation patterns or the expression of alternative splicing variants, which could generate products with different activity. We therefore set out to study the differential expression of NUMB splicing isoforms to determine whether these could be responsible for the observed variability in its function across the different databases and if NUMB could be proposed as a potential isoform-specific biomarker. According to the literature, only four NUMB transcripts generate protein-coding isoforms, those involving alternative splicing of exons 3 and 9. The sequences of these isoforms were compared with those of NUMBL, a close homologue of NUMB (**Figure 1G**). The longest isoform, p72, includes both exons. Loss of exon 3 results in the p71 isoform, while loss of exon 9 generates p66. Loss of both exons yields p65, the shortest isoform, which is also the most similar to NUMBL, which also lacks the regions homologous to exons 3 and 9 of NUMB [29, 47–49].

### A shift in NUMB isoform expression profiles characterises tumour samples

To analyse what happens in the expression of NUMB isoforms between non-tumour and tumour tissues, we selected the TCGA-BRCA dataset, choosing only patients with paired samples, to reduce noise due to the intrinsic variance of gene expression between individuals. TCGA-BRCA was selected because it provides the largest cohort of solid tissue normal samples paired with primary tumours, enabling robust intra-individual tumour–normal comparisons, which are not feasible in most other TCGA tumour types due to limited or absent matched controls. Firstly, we verified that the expression profile of each isoform did not vary significantly between the overall population and the paired samples, neither in the non-tumour condition nor in the tumour samples, so the paired samples constitute, for our study, a solid starting point (**Supplementary Figure 3**). Focusing on our results with paired samples, we observed two isoforms, p65 and p66, whose expression was reduced from non-tumour to tumour samples, while the other two NUMB isoforms, p71 and p72, clearly increased their expression in tumours. NUMBL, however, exhibited no changes in expression between non-tumour and tumour samples, suggesting that it is not affected by the oncogenic switch (**Figure 2A**). In addition, to ensure that this effect occurred intra-individually, we calculated the difference in the expression of each isoform for each patient between “tumour vs non-tumour”, showing that, in general, the expression of both p65 and p66 clearly decreased, while for the other two isoforms, p72 and p71, there is, again, a clear increase (**Figure 2B**). NUMBL, with three peaks, one centred on zero (with no changes), and two groups of samples with clear downregulation and clear upregulation, suggests the involvement of different regulatory events (**Figure 2B**).

**Figure 2.**
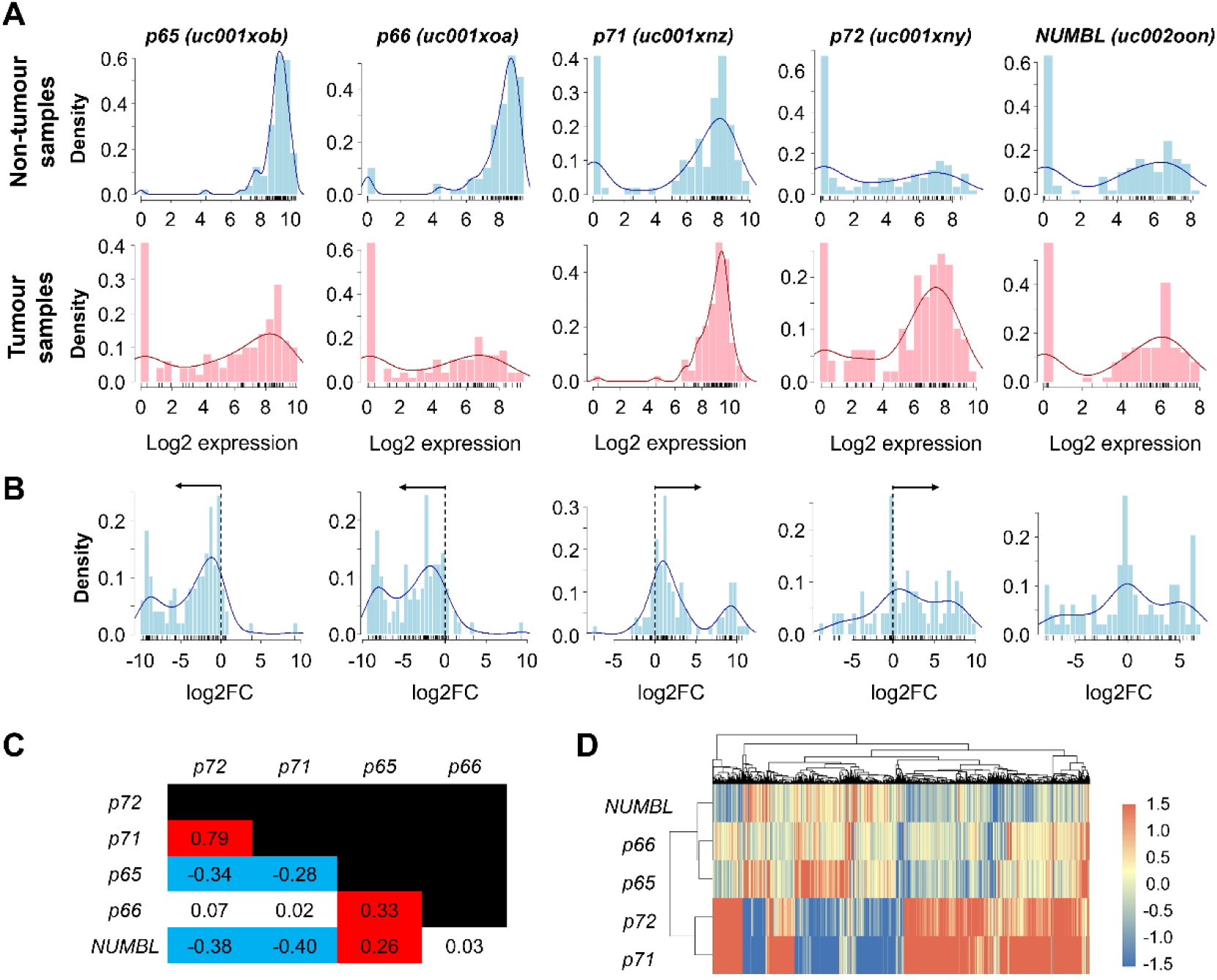
Tumour-associated changes in NUMB and NUMBL isoform expression and coordinated isoform behaviour. **A)** Expression density plots show a shift in NUMB isoform usage between paired tumour and non-tumour samples. Specifically, expression of p65 and p66 isoforms is reduced in a subset of tumour samples, whereas expression of p71 and p72 is increased. Expression of NUMBL appears relatively stable between conditions. These patterns suggest a tumour-associated shift in NUMB alternative splicing, favouring isoforms potentially linked to protumorigenic signalling. **B)** The distribution of paired log2-FC (tumour vs non-tumour samples) between NUMB isoforms highlights a coordinated shift in isoform usage. The p65 and p66 isoforms show predominant downregulation in tumours, whereas p71 and p72 show increased expression. In contrast, NUMBL shows a more balanced distribution, with both upregulated and downregulated cases in patients. This further supports a tumour-specific shift in NUMB isoform regulation, rather than a global suppression of NUMBL. Arrows indicate the predominant direction of expression change in tumour samples for each isoform. **C)** Correlation matrix showing the similarity between the differential expression profiles (log2FC) obtained by stratifying patients according to high or low expression of each NUMB isoform. The values represent Pearson’s correlation coefficients. Red indicates positive correlations, whereas blue indicates negative correlations. Only correlations with an absolute Pearson correlation coefficient ≥ 0.2 are highlighted, while correlations below this threshold are considered weak and therefore left uncoloured. **D)** Hierarchically clustered heatmap filtered by |log2FC| > 1.5 highlights a clear separation of gene expression patterns based on NUMB isoform stratification. Notably, isoforms p72 and p71 cluster together and show similar regulation patterns, while p65 and NUMBL also display correlated profiles. Opposing regulation blocks are evident between these two isoform groups, suggesting functionally divergent transcriptional programs associated with NUMB splicing. Columns represent differentially regulated transcripts/genes. Heatmap colour scale was limited to –1.5 to +1.5 for visualization purposes.

To further explore the implications of these results, we analysed the correlations of each isoform with all genes in the database, interpreting correlations in relative terms to identify convergent or divergent behaviour among NUMB isoforms rather than to formally test pairwise statistical significance. This yielded a correlation matrix that revealed a high similarity between p72 and p71, with a very high correlation (R=0.792). Similarly, we found positive correlations for both p66 and p65, as well as between p65 and NUMBL. This indicates some level of similarity, though with greater divergence, which could imply share regulatory effects on specific pathways, albeit through different mechanisms (**Figure 2C**). At the global level, NUMBL and p65 negatively correlated with p72 and p71, suggesting that these isoform groups may be involved in distinct regulatory programs. Considering the previously observed multimodal distribution of NUMBL, this correlation likely reflects coordinated regulation in a subset of patients, rather than a uniform pattern across the cohort. In contrast, the apparent lack of correlation of p66 with p72, p71, or NUMBL was striking, suggesting that this isoform may be regulated independently of the others.

Finally, we analysed the genes that correlated with each of the isoforms, based on the tumour vs. non-tumour expression differences (Δlog2FC) calculated for each gene in the paired samples. We applied a filter to select only those genes with an absolute Δlog2FC ≥ 1.5 for at least one of the isoforms analysed. These genes were used to construct a hierarchical heatmap representing their expression profiles, which revealed a clear separation between the p72/p71 isoforms and the p66/p65/NUMBL group (**Figure 2D**). Genes clustered according to the isoform group with which their expression was more strongly associated, indicating that p72 and p71 share similar regulatory effects, distinct from those of p65, p66, and NUMBL. This pattern suggests the presence of two opposing transcriptional programs linked to NUMB isoform usage.

### NUMB isoforms define distinct tumoral regulatory networks

To analyse how each NUMB isoform affects tumour gene expression, we identified genes showing either positive or negative correlations with each isoform. We found a significant number of genes whose expression patterns showed opposite correlations between the p72/p71 group and the p66/p65/NUMBL group (**Figure 3A**). These contrasting associations pointed to potential differences in NUMB-influenced signalling pathways. To explore this further, we examined the expression of genes involved in the SHH, Notch, Hippo, and WNT pathways, all of which are closely linked to tumour development and progression [31, 33, 47, 49–52]. NUMB is known to regulate Notch signalling through its interaction with NICD, which also connects functionally to the other considered pathways, SHH, Hippo and WNT. Our analysis revealed clear expression differences between the isoform groups, suggesting that each may be associated with distinct regulation patterns of these pathways depending on their prevalence in the tumour (**Figure 3B**). Thus, patients with higher p72/p71 transcriptional levels showed transcriptional profiles consistent with increased Notch/WNT/SHH pathway output and reduced Hippo pathway signalling. In the Notch pathway, we observed increased expression of genes such as *HES1*, *HEY1*, *PSEN2*, *JAG2*, and *DLL3*, along with reduced expression of *TLE1*, *TLE4*, *MFNG*, and *NUMBL*. The WNT pathway showed a stronger activation-associated pattern, with both p72 and p71 sharing similar gene signatures, with upregulation of canonical pathway components (*CTNNB1*, *TCF7*, *TCF7L2*, *LEF1*, *LGR6*, *WNT7B*, *FZD7*, *MMP7*, and others) as well as non-canonical ones (*WNT5A*, *ROR2*, *MAP3K7*, *CAMK2B*). Conversely, we observed marked downregulation of WNT inhibitors or genes typically described as suppressed upon pathway activation, such as *SFRP4*, *DKK2*, *SIAH1*, *CAMK2D*, and *CAMK2G*. For the SHH pathway, patients with higher p72/p71 levels showed increased expression of *SMO*, *SCUBE2*, and *HHAT*, all consistent with increased pathway activity. At the same time, we detected a clear decrease in *SUFU*, *PRKACB*, *SPOP*, and *CDON*, further supporting this interpretation. Finally, in the Hippo pathway, there was a notable increase in *YAP1*, *TEAD2*, *WWC1*, and *WWTR1*, together with reduced expression of *YWHAZ*, *YWHAE*, *YWHAG*, and *LATS2*, consistent with reduced Hippo pathway signalling and increased expression of YAP/TAZ-associated targets. On the other hand, patients with higher expression levels of the p66, p65, and NUMBL isoforms, displayed a more heterogeneous profile or reduced expression of the genes previously associated with pathway activation. We observed lower transcriptional levels of key components from the Notch pathway (*HES1*, *PSEN2*, *DLL3*), the WNT pathway (*TCF7L2*, *CTNNB1*, *WNT7B*, *LGR6*), and the SHH pathway (*SMO*, *SCUBE2*, *HHAT*). In contrast, the Hippo pathway showed a clearer pattern of increased pathway activity, particularly for p65, as reflected by decreased expression of *YAP1* and increased levels of *LATS2* and other upstream regulators.

**Figure 3.**
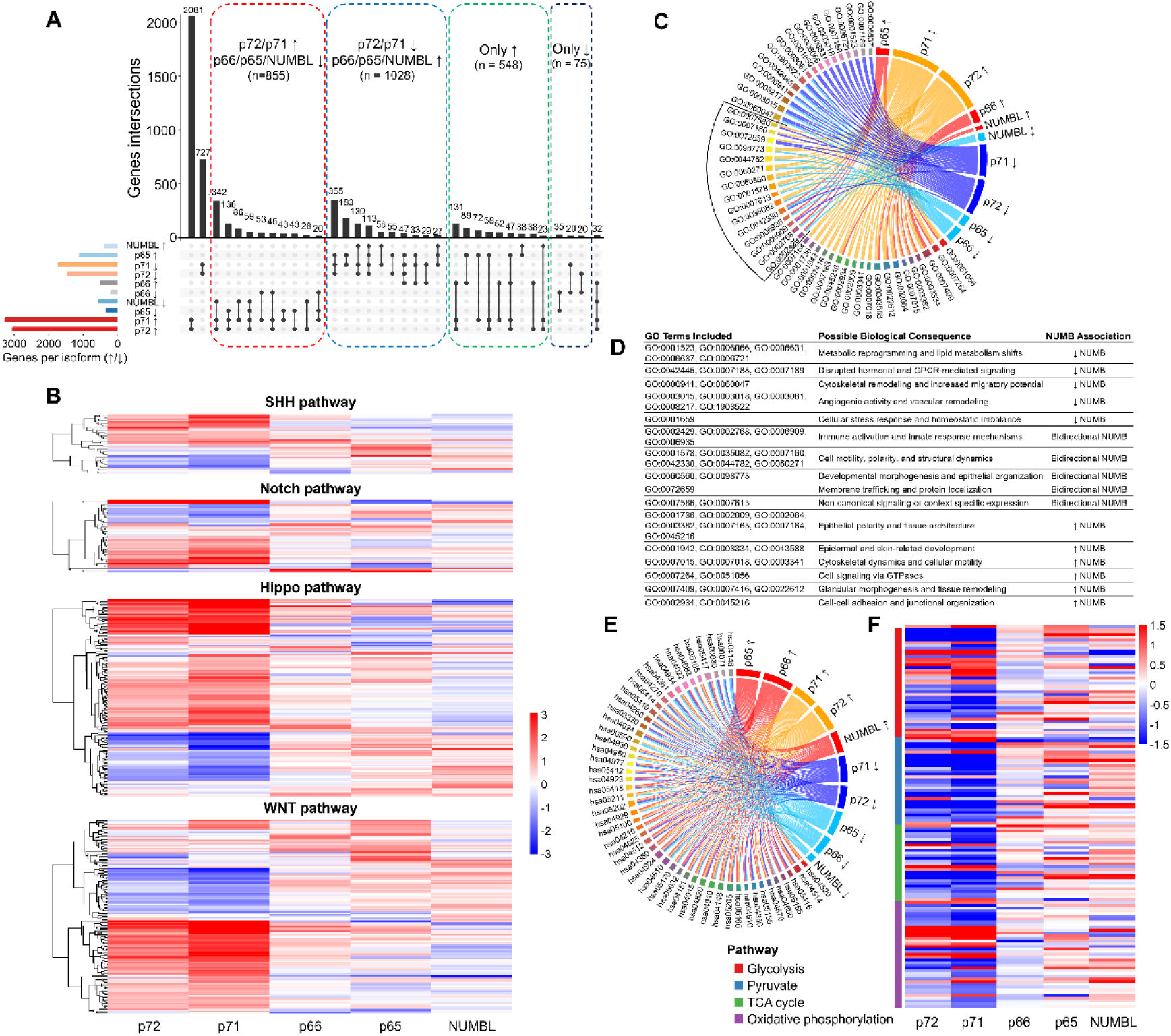
Isoform-specific functional associations of NUMB and NUMBL in tumour samples. **A)** UpSet plot illustrating the shared gene associations across NUMB isoforms. Distinct regulatory patterns emerge, separating p72/p71-up and p66/p65/NUMBL-down profiles from the inverse configuration. Categories were manually grouped by functional behaviour. Genes were included if they showed a log2FC > 1 and a p-value < 0.05 in at least one of the five isoforms, meaning not all genes are significantly differentially expressed across all conditions. Given the expected functional redundancy among some isoforms, the absence of significant expression changes in some of them is not unexpected. **B)** Non-hierarchical heatmaps illustrating the relationship of each NUMB isoform with signalling pathways previously associated with NUMB. Rows represent genes within each pathway (SHH, Notch, Hippo, and WNT), while columns represent samples ordered by condition or isoform correlation. Only genes showing an absolute log_2_ fold change ≥ 0.8 in at least one NUMB isoform were included in the input dataset. **C)** Chord diagram showing Gene Ontology (GO) terms enriched in genes associated with upregulated (↑) or downregulated (↓) NUMB isoforms. GO terms are grouped by regulation type: downregulated (top), bidirectional (middle, black-bordered), and upregulated (bottom). Opposing patterns between p72/p71 and p65/p66/NUMBL highlight divergent functional roles linked to NUMB isoform shifts. **D)** GO terms from panel C were grouped by biological function to interpret the biological impact of NUMB isoform-specific regulation. Terms classified as “bidirectional NUMB” show opposite associations between isoforms (e.g., upregulated in p72/p71 and downregulated in p65/p66/NUMBL), suggesting that alternative splicing of NUMB drives isoform-specific engagement of distinct cellular programs. In contrast, terms linked exclusively to either up- or downregulated isoforms may reflect functions that are shared or commonly regulated among isoforms with similar activity. **E)** KEGG pathway enrichment showing widespread associations between NUMB isoforms and multiple cellular pathways. Both upregulated and downregulated isoforms are linked to the same pathways, suggesting that different isoforms may modulate the same biological processes in opposing directions. This functional overlap supports the idea of isoform-specific regulatory programs. **F)** Heatmap showing the expression of genes from key metabolic pathways associated with NUMB isoforms. p72 and p71 show a coordinated upregulation of glycolytic genes and downregulation of mitochondrial respiration pathways, suggesting a shift toward aerobic glycolysis (Warburg effect). In contrast, p65, p66, and NUMBL maintain or promote expression of mitochondrial genes, highlighting functional divergence in metabolic regulation between NUMB isoforms.

To better understand the biological implications of these expression patterns, we performed a GO term analysis represented by a chord diagram (**Figure 3C**). GO terms were classified into three categories: (i) terms linked to positive correlations with isoforms from either the p72/p71 or p66/p65/NUMBL groups; (ii) terms associated with negative correlations; and (iii) a mixed group, where genes showed opposing correlations between the two isoform sets. In this last group, we identified functional categories related to immune response, cell motility, polarity, membrane trafficking, and signalling modulation (**Figure 3D**).

Many of these processes have been previously associated with NUMB functions, suggesting that isoform expression patterns may be associated with distinct functional biases in tumours. To further assess the correspondence between the different isoforms, we compared the Normalized Enrichment Scores (NES) obtained from gene set enrichment analysis (GSEA) (**Supplementary Figure 4**). This analysis revealed strong correlations between the isoform pairs p72/p71, p66/p65, p66/NUMBL, and p65/NUMBL, with gene sets consistently enriched (either positively or negatively) in the same direction for both isoforms, indicating convergence at the level of pathway-associated gene sets rather than identical functional outcomes. In contrast, p72 and NUMBL displayed complete opposition in their NES profiles, providing clear evidence of antagonistic regulatory biases in the modulation of cellular pathways. A similar pattern was observed for p71 and NUMBL, with only two gene sets enriched in the same direction, and for p72/p71 versus p65, where an opposing trend dominated, although a moderate number of gene sets still showed matching signs. Finally, the comparison between p72/p71 and p66 revealed a more heterogeneous distribution of NES values, suggesting potential functional divergence or a mixed regulatory behaviour.

We then created a second chord diagram using KEGG pathway annotations which revealed a more complex and interconnected landscape, reflecting potential cross-talk between pathways modulated by the different NUMB isoforms (**Figure 3E**). Some of the most frequent KEGG terms were related to intracellular signalling, immune response or metabolism. We performed a GSEA analysis for some of the KEGG terms found, observing a clear association between the p72/p71 pair terms, on the one hand, and the p66/p65/NUMBL terms, on the other (**Supplementary Figure 5A**), which suggests an opposite regulation in key processes. Thus, we observed a clear negative relationship of p72/p71 isoforms with mitochondrial processes, such as “*2-oxocarboxylic acid metabolism*” (hsa01210), “*alanine; aspartate and glutamate metabolism*” (hsa00250), “*fatty acid degradation/metabolism*” (hsa00071, hsa01212), “*glyoxylate and dicarboxylate metabolism*” (hsa00630) or “*citrate cycle (TCA cycle)*” (hsa00020), in this case, with an opposite relationship to that shown by NUMBL (**Supplementary Figure 5B**). This indicates that the expression of one type or another of isoforms is related to a less mitochondrial metabolism (p72/p71) or more based on OxPHOS (NUMBL). On the other hand, we also observed an opposite relationship between p72/p71 and p65/NUMBL isoforms at the level of “*cytokine-cytokine receptor interaction*” (hsa04060) or “*hormone signalling*” (hsa04081) (**Supplementary Figure 5C, 5D**), suggesting a differential regulation of immune and endocrine signalling pathways depending on the isoform expressed, which could impact tumour-stroma communication, immune infiltration, or hormonal responsiveness. Finally, we find terms exclusive to the p66/p65/NUMBL isoforms, such as “*Th1 and Th2 cell differentiation*” (hsa04658) (**Supplementary Figure 5E**), which is commonly associated with immune activation, suggesting a role for these isoforms in promoting anti-tumour immunity, in line with their previously proposed tumour-suppressive function [31, 32, 38, 41, 43, 51]. Similarly, other terms exclusively related to p72/p71, such as “*AMPK signalling pathway*” (hsa04152), whose negative NES value is consistent with its well-described tumour suppressor role through metabolic and immune regulation, thus supporting the protumorigenic role of the p72/p71 isoforms. Additionally, one the KEGG terms obtained was “*WNT signalling pathway*” (hsa04310), only for p71 isoform, being this finding consistent with our previous observation of increased correlation of p71 isoform with genes from this pathway. Together, these data reinforce the broader functional divergence observed between NUMB isoform groups. This dichotomy could reflect fundamental differences in the metabolic, immunological, and migratory behaviours of breast cancer cells depending on NUMB isoform usage, potentially linking specific isoform expression patterns to tumour aggressiveness or treatment response. We found a high number of metabolic processes connected to NUMB isoforms, including diverse categories such as amino acid, lipid, or carbohydrate metabolism. To better understand how these isoforms influence tumour bioenergetics, we analysed the expression of genes involved in glycolysis, pyruvate metabolism, the TCA cycle, and oxidative phosphorylation (**Figure 3F**).

Consistent with the GSEA findings, tumours enriched with p72/p71 isoforms showed negative correlations with key components of oxidative metabolism, consistent with reduced mitochondrial oxidative metabolic signatures. These isoforms were predominantly associated with strong negative correlations across key mitochondrial metabolic pathways, including core enzymes of the TCA cycle (e.g., *ACO1*, *CS*, *OGDH*, *MDH1*) and oxidative phosphorylation (e.g., *COX5A*, *ATP5MC3*, *NDUFB5*). Interestingly, a subset of mitochondrial genes (e.g., *ATP6V1B1*, *COX6B2*, *ATP12A*) displayed significant positive associations with p71, suggesting possible isoform-specific roles in mitochondrial remodelling rather than complete repression. In contrast, glycolytic and pyruvate-related genes such as *PKM*, *ENO2*, *GAPDHS*, and *LDHA* showed positive correlations with p72/p71, compatible with a possible metabolic reprogramming toward aerobic glycolysis, typical of the Warburg effect. These patterns are clearly represented in the heatmap (**Figure 3F**). Conversely, tumours enriched in NUMB isoforms p66, p65, or NUMBL presented more heterogeneous profiles, with frequent positive correlations across mitochondrial pathways and the TCA cycle, suggesting higher oxidative metabolism and bioenergetic flexibility.

### Dynamic remodelling of PDX tumours driven by NUMB isoform expression

Having confirmed the central role that changes in isoform expression can play in the molecular and functional evolution of tumours, we decided to analyse this evolution, taking advantage of the data available at the PDMR, which allows us to study isoform expression both in the original tumours and across the different tumour passages in mice. In this way, we were able to verify how both the p66 and p65 isoforms, more closely associated with a less tumorigenic phenotype and with a greater role for mitochondrial metabolism, as shown in the preceding section, significantly reduced their expression in the first PDX generated in mice (passage 0), without recovering their expression across passages (**Figure 4A**). However, we also found that there was a constant increase in the expression of the p72 and p71 isoforms, the most tumorigenic and with a more glycolytic metabolism, throughout the passages. Interestingly, a consistent increase in NUMBL expression was also observed, which may be involved in compensatory processes or in transcriptome stabilization across passages. When analysing how genes from the signalling pathways considered above correlated with each NUMB isoform, we observed a highly significant change from the original tumour (R) to passage 0, with multiple changes in gene expression in the four pathways, suggesting a global reorganization of the expression profile of these pathways, probably a consequence of molecular reprogramming or clonal selection after tumour implantation in the mouse (**Figure 4B**). Gene expression analysis of the four signalling pathways revealed a general downregulation upon tumour implantation in mice, followed by a progressive reactivation over subsequent passages (**Supplementary Figure 6A**). Notably, while the expression of many pathway-associated genes showed signs of recovery (**Supplementary Figure 6B**), the expression levels of the p66 and p65 isoforms remained consistently low (**Figure 4A**), suggesting a sustained suppression of the isoform profile associated with a less tumorigenic phenotype. All these changes point to a rewiring of regulatory relationships that likely reflect early reprogramming or selective pressures imposed by the host environment. To further dissect these dynamics, we applied dimensionality reduction using UMAP to pathway-specific gene expression profiles, uncovering the appearance of 5 to 8 distinct tumour clusters depending on the pathway considered (**Figure 4C**). For instance, tumours from the resection stage were largely confined to a single cluster in the Notch pathway, whereas SHH and Hippo pathways already exhibited broader initial heterogeneity. In the Hippo pathway, the formation of up to eight clusters suggests a particularly high degree of transcriptional plasticity and pathway-specific remodelling over time (**Figure 4D**). To better capture the temporal dynamics of these clusters and how tumours redistributed across them with each passage, we generated an Alluvial plot, which revealed both conserved and divergent transcriptional trajectories, further supporting the notion of progressive and pathway-specific remodelling over time (**Figure 4E**). These transitions revealed distinct evolutionary trajectories across pathways, with some clusters showing high conservation of tumour composition and others undergoing marked shifts, consistent with differential transcriptional plasticity and selective pressures acting on each pathway.

**Figure 4.**
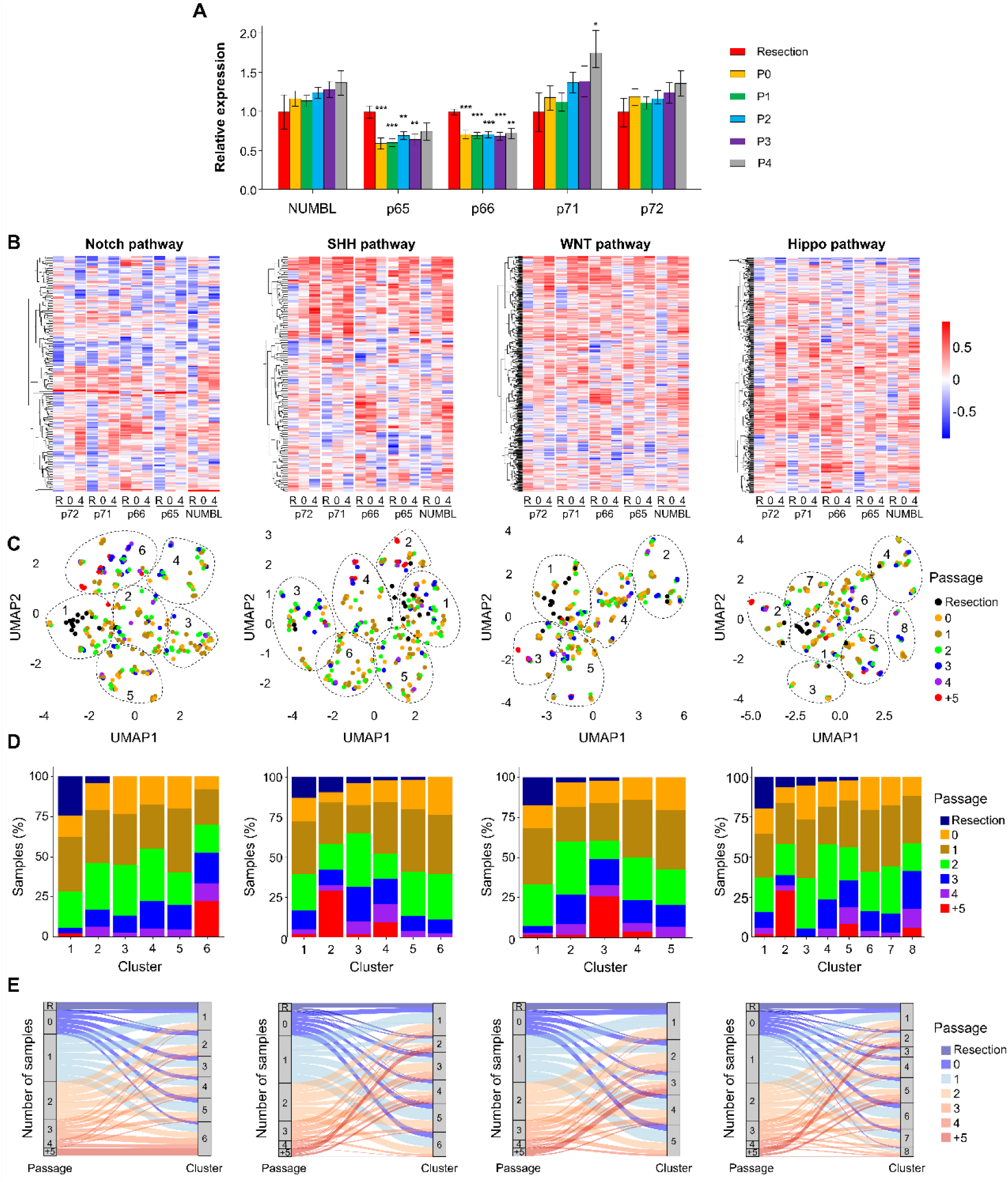
NUMB isoform dynamics drive pathway-specific transcriptomic reprogramming in patient-derived xenografts. **A)** Expression of individual NUMB isoforms and NUMBL across serial PDX passages (resection, P0–P4). A clear shift is observed: isoforms p65 and p66 are downregulated after implantation, while p72, p72 and NUMBL increase with subsequent passages. Statistical comparisons were performed between each passage and the resection baseline (*: p<0.05, **: p<0.01, ***: p<0.001). **B)** Correlation heatmaps between NUMB isoforms and the gene expression profiles of Notch, SHH, WNT, and Hippo signalling pathways. Heatmaps represent Pearson correlation coefficients (red = positive; blue = negative) and reveal distinct isoform-specific regulatory patterns for each pathway. **C)** UMAP-based clustering of tumour samples using pathway-specific gene sets reveals discrete clusters corresponding to transcriptional states. **D)** Bar plots showing the distribution of tumour passages within each cluster, indicating progressive transcriptional shifts associated with PDX passage. **E)** Alluvial plots depicting dynamic transitions of tumour samples between clusters across passages, highlighting the progressive reprogramming of signalling pathways over time.

### NUMB Isoform balance predicts signalling pathway activity and drug response in breast cancer cell lines

After confirming the progressive evolution of tumours in the PDX model, we decided to study what was happening with another of the most widely used models in laboratories: cell lines. To do so, we turned to the breast cell lines available at the CCLE, thus obtaining a list of 51 breast cell lines, 50 tumour cells from different origins, and the immortalized HMEL cell line. Since in this case we did not have a control group, as in the case of breast tumours, nor a starting point, as in the PDMR, we decided to use another approach to separate the cell lines based on some inherent property of the NUMB isoforms. Thus, we defined the *NUMB-score* based on the difference between the average expression of the p72/p71 isoforms minus the average expression of the p66/p65/NUMBL isoforms. Although this is an indirect approximation, this score allowed us to stratify the cell lines based on the relative predominance of potentially tumorigenic isoforms versus suppressive ones. To further separate the two groups, we eliminated from the comparison the 10% of the lines with the most intermediate values, thus obtaining a cutoff of around 0.5 (**Figure 5A**) and reducing the comparison to two groups of 23 cell lines.

**Figure 5.**
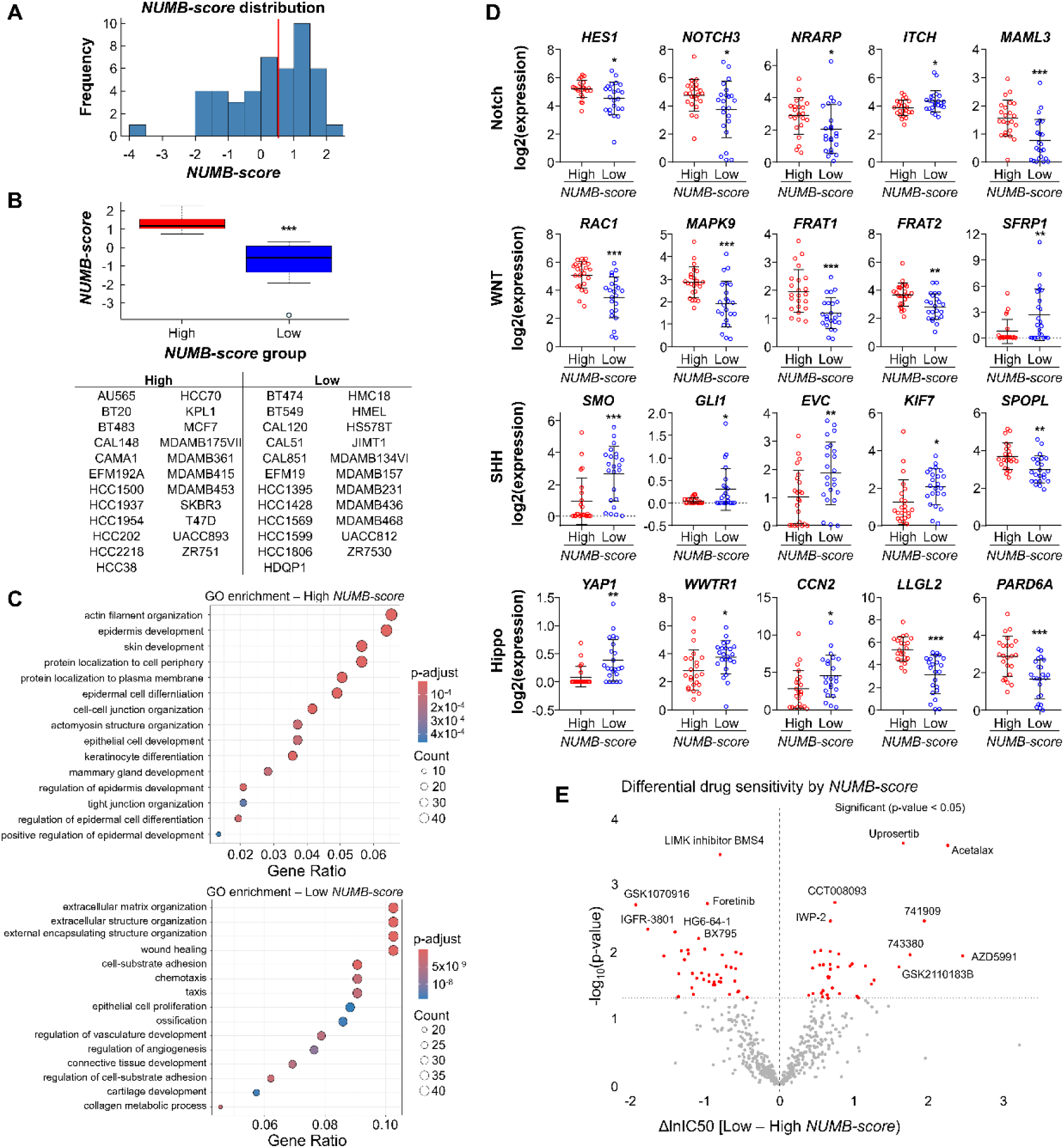
Functional analysis of the *NUMB-score* in breast cancer cell lines from the CCLE. **A)** Distribution of the *NUMB-score* in the 46 cell lines selected for the study (23 per group), with the cutoff point indicated by the red line. **B)** *NUMB-score* values for each group, with the list of cell lines included in the high and low groups. **C)** GO enrichment dot plots for both groups, showing greater enrichment in development and differentiation-related terms in the high *NUMB-score* group, and enrichment in extracellular matrix and motility-related terms in the low group. **D)** Expression of representative genes from the Notch, WNT, Hippo, and SHH pathways, illustrating increased activity of the first three pathways in the high *NUMB-score* group. Transcript identifiers used for each pathway component are provided in **Supplementary Table 4**. **E)** Volcano plot of drug sensitivity analysis; most compounds show higher efficacy in cell lines with a high *NUMB-score*.

Although the heterogeneity is obviously greater and we cannot consider greater or lesser tumorigenicity based on this system, since we were dealing with 50 tumour lines with thousands of passages, it was striking that the only non-tumour line, HMEL, fit with this system in the group with the lowest *NUMB-score*, which meant it fell into the group with the highest p66/p65/NUMBL ratio (**Figure 5B**). Once both group cell lines were separated, we could observe that they presented a different enrichment of GO terms. Thus, lines with a higher *NUMB-score* had a greater number of GO terms related to development and differentiation, as well as processes related to protein transport and cell attachment, while those with a lower *NUMB-score* had a higher proportion of terms associated with extracellular matrix remodelling, motility, or angiogenesis (**Figure 5C**). This apparent contradiction could be due to contextual functions of NUMB isoforms in cell lines highly adapted to culture, where invasive traits can emerge despite a higher proportion of suppressor isoforms, possibly due to selection pressure or reconfiguration of signalling networks. Although the low-score group shows a more invasive profile, the high *NUMB-score* is associated with greater epithelial organization and plasticity, which could reflect an early phase of high proliferative potential and tumour adaptation. Following this, we decided to examine what happened to key genes of the different signalling pathways analysed throughout the study. We observed clear activation of the Notch pathway in the group with the highest *NUMB-score*, as shown by the activation of genes such as *HES1*, *NOTCH3*, *NRARP*, and *MAML3*, and the inhibition of *ITCH* (**Figure 5D**). Similarly, we also observed that the WNT pathway was more active in the high *NUMB-score* group, with greater activation of the genes *RAC1*, *MAPK9*, *FRAT1*, and *FRAT2*, and less activation of the gene *SFRP1*. The SHH pathway, on the other hand, appeared more activated in the group with the lowest *NUMB-score*, with this group having more activity of the genes *SMO*, *GLI1*, *EVC*, and *KIF7*, and less activity of *SPOPL*. Finally, the Hippo pathway also appeared more activated in cells with the highest *NUMB-score*, with lower expression levels of *YAP1*, *WWTR1*, and *CCN2*, and higher expression levels of *LLGL2* and *PARD6A*. These results indicate that the balance of NUMB isoforms could modulate the functional status of key pathways associated with tumour proliferation, differentiation, and plasticity, with distinct activation profiles depending on the predominance of one isoform or another. Having confirmed the differences at the phenotypic level and in the activation of signalling pathways, we decided to test whether there were differences in drug resistance between the cell lines. To do this, we used the IC50 data available at the GDSC and performed a Volcano plot separating the cell lines based on the *NUMB-score*. We identified 78 drugs with significant differential sensitivity between the two groups, with 38 preferentially affecting high NUMB-score cells and 40 low *NUMB-score* cells (**Figure 5E, Supplementary Table 6**). These results indicate that the NUMB-score captures distinct drug-response profiles and supports its potential utility as a predictive marker of therapeutic sensitivity.

### Association of the NUMB-score with tumour features and survival in TCGA-BRCA

Survival stratification based on individual NUMB isoforms or NUMBL in the TCGA-BRCA cohort consistently resulted in markedly asymmetric patient groupings, reflecting cutpoints optimized toward expression extremes (**Supplementary Figure 7A**). Although p66 and NUMBL showed statistically significant associations with overall survival, these effects were observed only under highly unbalanced stratifications, whereas p72, p71, and p65 failed to reach significance across robustness analyses. Notably, p72 did not show prognostic relevance despite near-balanced group sizes, indicating that lack of association was not solely driven by group imbalance. Collectively, these findings indicate that individual isoforms provide limited and unstable prognostic information when considered in isolation. Application of the previously defined NUMB-score resulted in a more symmetric patient distribution and provided a more robust framework for clinical stratification, as illustrated by the NUMB-score density profiles in tumour and non-tumour samples (**Supplementary Figure 7B**).

Having established the relevance of the *NUMB-score* in cell lines, we sought to assess whether this metric could also be applied to TCGA-BRCA tumours by analysing the full cohort. We observed a general increase in its value in tumour samples compared to non-tumour tissues (**Figure 6A**). When stratifying tumour samples by stage, the score remained consistently elevated across all stages (**Figure 6B**), suggesting that the isoform switch is an early event in tumour evolution and is maintained throughout progression. Using the AJCC classification, we found that although all tumour subtypes exhibited higher *NUMB-score* than non-tumour tissues, Her2+ tumours had the highest values (**Figure 6C**). Stratification of patients into high and low NUMB-score groups (**Figure 6D**) revealed a significant difference in overall survival when considering the full cohort (**Supplementary Figure 7C**). Importantly, reducing the influence of a small fraction of highly influential samples (∼1% of the cohort) further strengthened the separation between groups while preserving comparable group sizes (**Figure 6E**), supporting the robustness of the *NUMB-score* as a prognostic indicator. Together, these results show that integrating isoform information into a composite score provides substantially improved clinical stratification compared with analyses based on individual NUMB or NUMBL isoforms.

**Figure 6.**
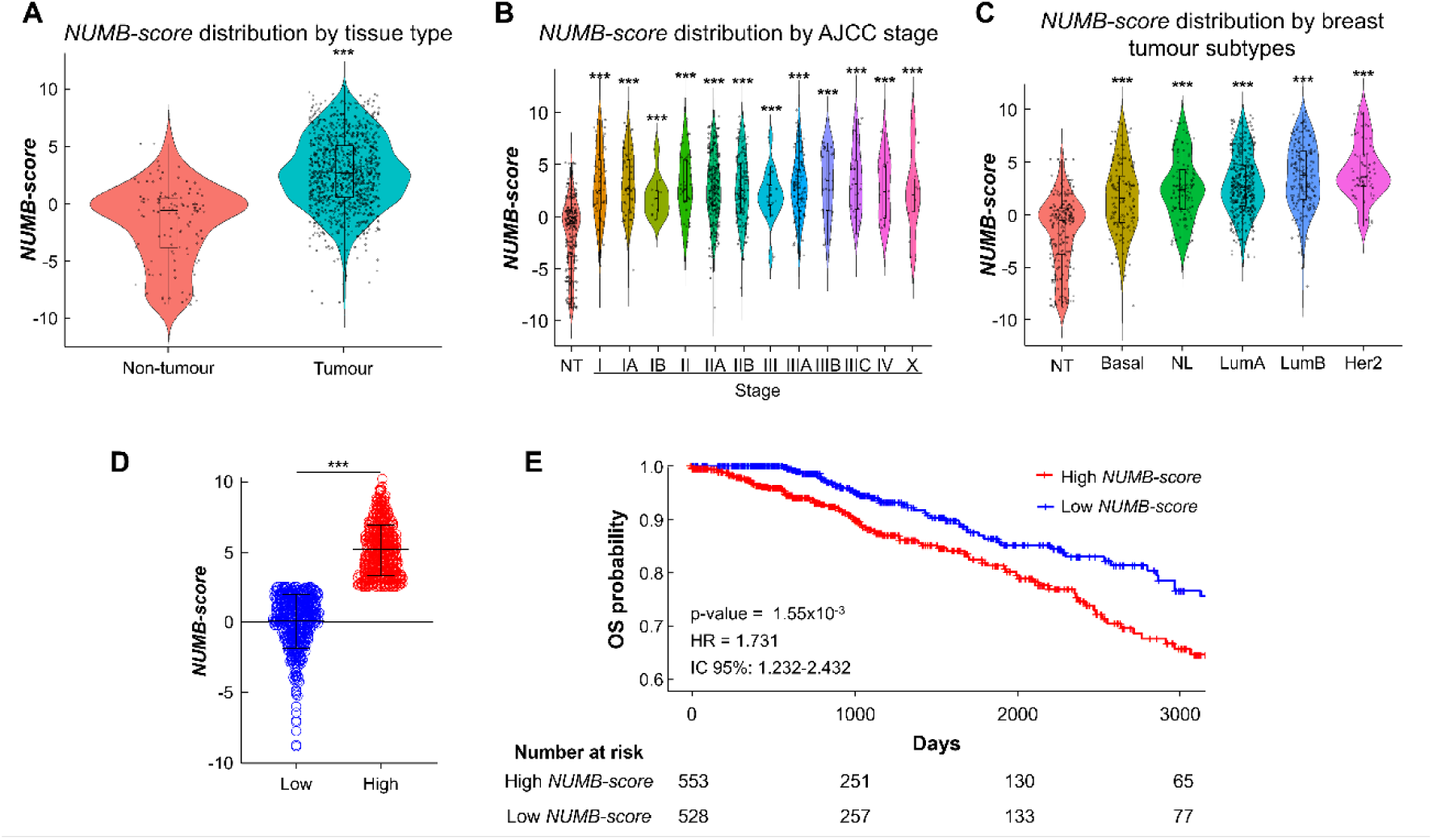
High *NUMB-score* is associated with breast tumours and poorer overall survival. **A)** *NUMB-score* is significantly higher in tumour versus non-tumour tissues. **B)** Elevated *NUMB-scores* are consistent across all AJCC tumour stages. **C)** All breast cancer subtypes show increased *NUMB-scores*, with Her2+ tumours having the highest levels. **D)** Tumours were stratified into low and high NUMB-score groups, showing a significant separation. **E)** Kaplan–Meier survival analysis indicates worse overall survival in the high NUMB-score group. ***: p-value < 0.001.

## DISCUSSION

NUMB is widely studied in development and differentiation as a key regulator of pathways such as Notch, WNT, SHH, and Hippo [27, 31, 36, 38, 53]. These pathways have long been known to show aberrant activity in a large percentage of tumours, regardless of their location, making it essential to gain a deeper understanding of the role played by NUMB. However, previous studies have reached divergent conclusions on whether NUMB acts as a tumour suppressor or an oncogene [28, 43, 44, 51, 54, 55], motivating an in-depth analysis of potential alterations in its expression. NUMB generates multiple isoforms through AS, mainly via inclusion or exclusion of exons 3 and 9, which markedly affets its functional behaviour. While exon 3 status can modulate NUMB interactions with other proteins [29, 53, 56], it does not seem to be critical for regulation of the signalling pathways analysed in our study, as most alterations are linked to exon 9 inclusion/exclusion. Transcriptomic data show that exon 9-containing p72/p71 isoforms are strongly associated with activation of these pathways, in turn relating to uncontrolled proliferation, evasion of differentiation, and tumour microenvironment remodelling. Both isoforms appear to be associated with metabolic reprogramming, with less dependence on mitochondrial metabolism and, notably, increased sensitivity to certain antitumour treatments. NUMB has previously been linked to mitochondrial metabolism; as its depletion enhances glycolysis and promotes mitochondrial fragmentation, although these effects were not assigned to specific isoforms [57, 58]. Overall, this supports a broader functional reprogramming beyond the signalling pathway alterations analysed. In contrast to p72/p71, exon 9-null p66/p65 isoforms are associated with transcriptional programs enriched in inflammation, immune response and epithelial differentiation, and show a stronger link to oxidative phosphorylation, consistent with active mitochondrial metabolism, typical of healthy tissue and a less aggressive phenotype. Together with previous research, our data indicate that both isoform groups are functionally active but drive different cellular programs [47, 55, 59, 60], with p72/p71 promoting a more aggressive tumour phenotype and p66/p65 correlating with reduced proliferation and potential antitumour properties. In fact, and according to our results, increased p72/p71 expression is observed not only in breast cancer but also in other tumours (e.g.,lung LUAD/LUSC and medulloblastoma) [55, 61], suggesting that NUMB AS is a common mechanism favouring expression of higher oncogenic potential isoforms, as we observed in both TCGA-BRCA and PDX samples. Conversely, higher expression of exon 9-null isoforms in normal tissue may help maintain the balance between proliferation and differentiation. Overall, our findings show that splicing alterations, already recognized as a hallmark of cancer [3, 14, 16], can be used as a molecular stratification tool to enable isoform-based personalized medicine strategies. These results indicate that NUMB’s function is isoform-specific and context-driven rather than that of a general tumour suppressor, and that the isoforms involved may exert opposing effects: protective in physiological conditions but potentially oncogenic in tumour contexts. Differential NUMBL expression strongly correlated with the p65 isoform, suggesting that NUMBL could behave as a functional “fifth isoform” of NUMB, especially in contexts where tumour-suppressive activity is maintained. In our analysis, NUMBL levels were similar in healthy and tumour tissues, supporting independent regulation and indicating that NUMBL does not compensate for NUMB isoform switching. The NUMB isoform switch was consistently observed across TCGA transcriptomic data, in vivo PDX models and cell lines, supporting the robustness of this phenomenon. Previous studies report that NUMBL loss confers resistance to multiple chemotherapeutic agents in breast and cervical cancer cell lines, accompanied by Notch activation and cancer stem cell markers [62]. In addition, exon 9 inclusion is linked to Notch activation in non-small cell lung cancer [63], and loss of p66/p65 correlates with resistance to genotoxic agents in breast carcinoma [64]. Our data confirm and extend these findings by showing that (i) the same switch occurs in breast PDX, (ii) it correlates with tumour status in TCGA-BRCA, and (iii) the NUMB-score predicts chemotherapy response and overall survival. Notably, breast cancer PDX analyses also indicate a phenotypic shift following implantation, with tumours initially enriched for suppressor isoforms (p66/p65) frequently switching toward tumour-associated isoforms (p72/p71), suggesting in vivo selection pressures that favour clones expressing oncogenic NUMB isoforms and supporting NUMB as a dynamic regulator of tumour plasticity. The rapid loss of p66/p65 and progressive gain of p72/p71 post-implantation appoint to selection for “isoform-specific plasticity”, consistent with evidence that long NUMB isoforms (p72) promote recycling of proliferative receptors (e.g., ALK membrane relocalization) to sustain signalling and confer adaptive advantages in challenging microenvironments [59]. Our longitudinal follow-up supports that this switch is not a culture artifact but a recurrent in vivo adaptation route, implying that NUMB isoforms dynamics may insight into tumour status and clinical evolution, including progression and therapy resistance.

NUMB has traditionally been considered a tumour suppressor [31, 32, 38, 41, 43, 51], due to its ability to regulate multiple signalling pathways, including those examined in our study, but its impact appears to depend on the predominantly expressed isoform. Evidence from cancer cell lines (e.g.MCF-7) [41], and tumour samples [54, 55, 61, 65] links exon 9–containing isoforms (p72/p71) to more aggressive phenotypes and increased proliferative activity, partly through modulation of key components of the AKT and WNT pathways that promote tumour growth [31, 53, 65]. In contrast, exon 9-null isoforms are more common in differentiated or non-tumour cells and are linked to less aggressive behaviour and, in some cases, a more stable epithelial state [59, 61, 66]. Our analyses further suggest they have limited capacity to modulate pro-proliferative pathways such as Notch or WNT. This bimodal behaviour explains why studies that only measure global NUMB expression have reached contradictory prognostic conclusions.

Breast cancer provided a clear model to study how shifts in NUMB isoform expression toward to exon 9-containing variants can activate transcriptional programs that promote tumour progression. Rather than total NUMB loss driving tumour development, our results support that an imbalance in the relative abundance of NUMB isoforms is a critical event. Exon 9 inclusion in NUMB should not be viewed as intrinsically beneficial or detrimental, but as a feature that amplifies its functional capacity, which can lead to either anti- or pro-tumour effects depending on biological context (tissue type, the status of pathways such as Notch, WNT and SHH, and the tumour microenvironment). Our data support that NUMB does not have a single role in breast cancer; instead, its impact depends on the balance between isoforms, with exon 9-containing isoforms linked to stronger activation of Notch/SHH/WNT and Hippo repression, whereasp66/p65 correlate with inhibition of these same pathways. This suggests a genuine functional mechanism whereby exon 9 presence/absence reconfigures cellular signalling, challenging the classical view of NUMB as a static inhibitor of oncogenic pathways and supporting its role as a versatile adapter whose functionality depends on the predominant isoform and the specific cellular environment. Several mechanistic explanations can explain our findings. Exon 9 lies within NUMB’s PRR, which binds multiple SH3-domain proteins and modulates their interactions [67, 68]. In p72/p71 isoforms, this exon may support functional complexes that influence the trafficking, stability or transcription of Notch, WNT, and SHH receptors and effectors. Switching from p66/p65 to p72/p71 could shift NUMB from a pathway inhibitor to a facilitator, with p72/p71 acting as oncogenic scaffolding by recruiting regulators such as ITCH, aPKC, or other adapters to reshape signalling complexes. This may stabilize receptors (e.g. Frizzled in WNT or PTCH1 in SHH), promote NICD nuclear translocation or inhibit Hippo activators (e.g. LATS1/2), promoting YAP/TAZ cytoplasmic retention or degradation. Hippo repression is especially relevant, as its inactivation enables YAP/TAZ nuclear entry of YAP/TAZ and the transcription of growth-promoting genes [69]. In addition, there is extensive crosstalk: YAP can enhance WNT, suppress Notch-induced differentiation, and SHH may inhibit Hippo activity in specific contexts [70–72]. Therefore, p72/p71-mediated Hippo inhibition may indirectly reinforce Notch, WNT and SHH signalling, consolidating a more aggressive tumour phenotype.

Previous studies have suggested that AS-driven changes in PTB domain length could modulate intracellular NOTCH processing. Short-PTB isoforms (p65/p71) would favour endocytic retention and processing to NICD, facilitating nuclear entry [32, 59], whereas long-PTB isoforms (p66/p72) could restrict this step. Consistent with this model, lung cancer studies have described p66/p65 as Notch inhibitors andp72/p71 may enhance it, highlighting the functional relevance of isoform composition across tumour types [73]. NUMB is also a transcriptional target of canonical WNT/β-catenin pathway, which induces its expression [31, 74], potentially within feedback loops jn which NUMB modulates WNT in an isoform-dependent manner. In fact, our results suggest that p66/p65 inhibit WNT, whereas p72/p71 show an activating profile. Similarly, previous work linked NUMB loss to increased SHH activation [37, 51], although without isoform specificity. Our data refine this model by showing that SHH activation is associated only with specific NUMB isoforms.

Our earlier work proposed NUMB as an inhibitor of Notch, SHH, and WNT [47], but the present analysis refines this view by showing that this regulatory capacity is isoform-dependent. Specifically, p66/p65 and NUMBL display patterns consistent with inhibition of these pathways, whereas p72/p71 are associated with their activation, supporting the need to analyse NUMB as a set of functionally distinct isoforms. Mechanistically, MAPK/ERK signalling has been reported to promote exon 9 inclusion and favour p72/p71 expression in lung cancer cells, supported by PTBP1 and antagonized by ASF/SF2, while QKI-5 represses exon 9 inclusion by competing with SF1 at the branch point [28, 53, 75]. Extending these findings, we show in a cohort of more than 1,000 breast cancer patients that the shift from predominant p66/p65 expression to p72/p71 represents a robust and early transcriptomic signature of tumour status. This is in line with functional studies reporting that exon 9-inclusive isoforms enhance Notch signalling and increase target gene expression [28, 63]. Accordingly, p72/p71-enriched tumours exhibit a transcriptomic signature of Notch activation in both TCGA-BRCA and PDX models. Overall, these results support an isoform-dependent model that departs from the classical view of NUMB function: p72/p71 are associated with activation of Notch, SHH and WNT and repression of Hippo, whereas p66/p65 show the opposite pattern. This functional divergence in key pathway activity could directly influence cell fate and tumour behaviour.

## CONCLUSIONS

Beyond their biological relevance, these results support an emerging precision-medicine paradigm in which functional gene analysis should consider not only total expression but also isoform-specific contributions. Since splicing alterations enable tumour cells to fine-tune their proteome, identifying and quantifying NUMB isoforms provides a robust basis for patient stratification and targeted therapy design. In line with this, we developed the *NUMB-score*, an expression-based contrast metric capturing the balance between tumour promoter NUMB-isoforms (p72/p71) and the suppressive group (NUMB p66/p65 and NUMBL). The NUMB-score stratified cell lines into groups with divergent functional profiles and drug sensitivities, suggesting potential utility for predicting therapeutic responses. Translationally, we propose the NUMB-score as a simple metric to quantify isoform balance defined by exons 3 and 9, complementing prior work on tissue patterning and cancer stem cells [60, 66]. We further supported this approach by(i) validating it in over 50 CCLE lines, (ii) correlating the score with response to 18 drugs in GDSC, and (iii) associating it with overall survival in TCGA. Together, these findings support its clinical potential as a splicing-derived transcriptomic marker for more precise personalized diagnosis and treatment.

Our findings help reconcile the dual tumour-suppresive and oncogenic roles attributed to NUMB by showing that different isoforms can exert opposing functions. This finding has important implications for interpreting previous studies and for designing new therapeutic strategies, as treating NUMB as a single functional entity can lead to misinterpretations. More broadly, alterations in AS enable tumour cells to remodel their proteome by favouring isoforms linked to proliferation, migration, or immune evasion, and our results demonstrate that NUMB undergoes systematic isoform switching in the tumour context, with clear functional implications. Taken together, our data suggest that NUMB should not be considered a single marker gene, but rather a functional family of isoforms with specific properties, whose differential expression modulates key signalling pathways and could predict therapeutic response. Therefore, the study of AS not only provides biological knowledge but also has direct potential in precision medicine, both in diagnosis and in the stratification and prediction of drug response.

## METHODS

### Functional analysis of isoform switches

To obtain a quantitative summary of isoform switch events with functional consequences, the IsoformSwitchAnalyzeR package (RRID:SCR_027320) implemented in R (RRID:SCR_001905) via Bioconductor (RRID:SCR_006442) was used [76, 77]. From the switchAnalyzeRlist object containing the significant results of the isoform expression analysis between tumour samples and controls [76], tumour-type-specific subsets were generated for each TCGA tumour type (RRID:SCR_003193). Only events with functional consequences on the protein were selected (e.g., domain gain/loss, peptide signal, intron retention, among others). For each cancer type, the number of affected genes, the total number of isoform switches, and the number of isoforms involved were quantified. Additionally, global patterns of alternative splicing events were analysed across all tumour types using the analyzeAlternativeSplicing() function. Events were grouped by category (exon skipping, intron retention, alternative acceptor/donor sites, etc.) and summarized per cancer type. A separate analysis focused on breast cancer (BRCA) compared tumour and non-tumour samples by counting the number of AS events of each type detected in tumour (T) and non-tumour (N) samples.

### Web tools and public data resources

The cBioPortal for Cancer Genomics (RRID:SCR_014555) was used to explore potential alterations in NUMB that may contribute to tumorigenesis [45]. This platform provides gene-level alteration frequencies across multiple cancer types. In addition, differential expression of NUMB between 31 TCGA tumour types and their corresponding normal tissues (from the GTEx database, RRID:SCR_013042) was analysed using GEPIA2 (RRID:SCR_026154) [46], which also reports TPM values. To assess the prognostic relevance of NUMB expression in breast cancer, survival data were obtained from the R2 Genomics Analysis and Visualization Platform (RRID:SCR_025770). To reduce platform-related variability, only Affymetrix-based datasets were included (HG-U133A, GEO platform GPL96; HG-U133 Plus 2.0, GEO platform GPL570). NUMB expression was measured using probe 209073_s_at across publicly available datasets (GSE69031, GSE21653, GSE1456, GSE20685, GSE25066, GSE42568, GSE162228, and GSE12093) from the Gene Expression Omnibus (GEO, RRID:SCR_005012). [78–85]. This analysis aimed to evaluate the association between NUMB expression levels and patient survival outcomes.

### Data source and preprocessing

The isoform expression data used in this study, restricted to breast samples, came from three public sources: The Cancer Genome Atlas (TCGA) [86, 87], the Patient-Derived Models Repository (PDMR, https://pdmr.cancer.gov) [88], and the Cancer Cell Line Encyclopaedia (CCLE, RRID: SCR_013836) [89].

Patient data were obtained from the TCGA-BRCA study through the Firebrowse portal (RRID:SCR_026320), using pre-normalized expression files generated with RSEM (RRID:SCR_013027) at the isoform level (*Level_3_RSEM_isoforms_normalized_data.data*). Gene assignment was performed using HGNC nomenclature (RRID:SCR_002827). For patient-derived models, isoform-level quantification files in RSEM format *(.isoforms.results*) corresponding to breast tumour samples were obtained from the PDMR repository and processed to extract *transcript_id*, *gene_id*, and *expected_count* columns. The files were integrated into a single table using *transcript_id* and *gene_id* as common keys, and isoforms with zero expression across all samples were removed. For cell lines, transcriptomic data from the CCLE (*CCLE_RNAseq_rsem_transcripts_tpm_20180929.txt*) were used, containing TPM values estimated with RSEM. Only lines classified as derived from mammary tissue were included, and gene annotations were assigned using an Ensembl–HGNC correspondence table (Ensembl: RRID: SCR_002344). In all cases, expression values were transformed using log2(x + 1), incorporating a pseudocount to avoid undefined values in cases of zero expression.

For subsequent analysis, specific NUMB and NUMBL isoforms of interest were identified and selected from each dataset. For the TCGA-BRCA and PDMR data, transcript identifiers from the UCSC Genome Browser (RRID:SCR_005780) were used, namely uc001xob (p65), uc001xoa (p66), uc001xnz (p71), uc001xny (p72), and uc002oon (NUMBL). For the CCLE dataset, the corresponding Ensembl identifiers were used: ENST00000554546 (p65), ENST00000535282 (p66), ENST00000557597 (p71), ENST00000555238 (p72), and ENST00000252891 (NUMBL). These identifiers were used as keys to extract and compare expression levels across sets.

### Isoform expression analysis in TCGA-BRCA

In the TCGA-BRCA set, the expression distribution of each isoform was evaluated using histograms and density curves generated with the *density()* function in R, both in the entire tumour cohort and in the subcohort of paired samples (tumour and non-tumour tissue from the same patient). The presence of bimodal distribution in the expression of each isoform was then assessed using Hartigan’s dip test (*dip.test()*, *diptest* package, version 0.77-2, CRAN) [90]. In cases with significant evidence of bimodality (p < 0.05), tumour samples were classified into high- and low-expression groups using k-means clustering (k = 2), using centroids as cutoff points. For isoforms with unimodal distribution, tertile-based stratification was applied, defining the “low” and “high” groups based on the first and third tertiles of the distribution, and excluding samples from the intermediate tertile.

To study changes in isoform expression at the intraindividual level with greater resolution, pairwise differential expression values were calculated for each patient. These log2FC values were obtained as the difference between the transformed expression levels in the tumour sample and its corresponding non-tumour sample from the same individual. The result was a matrix of log2FC values by patient and isoform, which served as the basis for correlation and differential expression analyses. This strategy allowed for the identification of coordinated patterns of change between isoforms, such as compensatory increases or decreases, that would not be evident in aggregated expression analyses at the population level. To visualize the global differential expression patterns associated with each NUMB isoform, log2FC matrices were generated from paired samples. In a first analysis, only those genes whose expression showed an absolute change greater than 1.5 (|log2FC| > 1.5) in at least one of the conditions analysed were selected, with the aim of highlighting pronounced regulatory events. Subsequently, the regulation of genes belonging to key functional pathways in cancer and energy metabolism was explored. To this end, transcripts annotated in the WNT (hsa04310), Notch (hsa04330), SHH (hsa04340), Hippo (hsa04390), Glycolysis (hsa00010), pyruvate metabolism (hsa00620), citric acid/TCA cycle (hsa00020), and oxidative phosphorylation (hsa00190) pathways were selected according to the KEGG database and intersected with the previously filtered set of differentially expressed genes (|log2FC| ≥ 0.8 and p < 0.05 in at least one isoform). The resulting matrices were represented by heatmaps generated with the R package *pheatmap* (RRID: SCR_016418) [91], using a colour scale centred on zero and hierarchical clustering by both rows (genes) and columns (isoforms).

To identify differentially expressed genes in common between the different NUMB and NUMBL isoforms, an UpSet intersection graph was constructed using the *UpsetR* package (RRID:SCR_022731) [92]. Subsets of genes were defined whose expression presented an absolute magnitude of change |log2FC| > 1 in each isoform, considering both upregulation and downregulation. From these subsets, a binary matrix was generated indicating the presence or absence of each gene in each of the ten conditions (isoform combination and direction of change). Genes shared by at least two sets were retained, applying a threshold of a minimum of 20 genes per combination.

To study biological processes and molecular pathways potentially associated with the differentially expressed genes according to each NUMB and NUMBL isoform, functional enrichment analyses were performed using Gene Ontology (GO, RRID:SCR_002811, Biological Process category) and KEGG database annotations. In both cases, subsets of significantly regulated genes were defined (|log2FC| > 0.8 and p < 0.001 for GO; p < 0.01 for KEGG), differentiating between up- and down-regulation. HGNC identifiers were converted to ENTREZ IDs (RRID:SCR_016640) using the *bitr()* function of the *clusterProfiler* package (RRID: SCR_016884) [93], and analyses were performed with *enrichGO()* and *enrichKEGG()* respectively. For each group, terms or pathways with a significant p-value were retained, limiting the 100 most significant terms to the total when the total number exceeded this threshold. Subsequently, those terms or pathways shared by at least two groups were selected. The 40 most frequent GO terms and 50 KEGG pathways were visualized using chord diagrams generated with the *circlize* package (RRID:SCR_002141) [94]. In both visualizations, nodes represent isoforms (grouped by regulatory direction) and enriched terms, connected by lines that reflect significant functional associations. Furthermore, in the GO diagram, the terms were organized according to their functional association pattern (positive, negative or mixed), thus facilitating the biological interpretation of the results. In addition, an isoform-specific GSEA was performed on GO-BP terms to further assess coordinated transcriptional programs associated with each NUMB and NUMBL isoform. For this purpose, genes were ranked independently for each isoform according to log2FC values, collapsing multiple transcripts per gene by selecting the most extreme log2FC while preserving directionality. GSEA was conducted using the *gseGO()* function of the *clusterProfiler* package, with HGNC symbols as identifiers and BP ontology, retaining gene sets with a minimum size of 10 and a maximum size of 500. NES were obtained for each isoform, and GO terms detected in at least two isoforms were retained for comparative analysis.

A complementary GSEA was also performed using KEGG pathway annotations. For each isoform, genes were ranked by log2FC after collapsing multiple transcripts per gene to the most extreme value while preserving directionality, and HGNC symbols were converted to ENTREZ identifiers prior to analysis. KEGG GSEA was carried out using the *gseKEGG()* function of *clusterProfiler*, retaining pathways with sizes between 10 and 500 genes. NES were obtained independently for each isoform, and enrichment profiles were subsequently compared across isoforms. In addition, mean NES values were computed for two isoform groups (p72/p71 and p66/p65/NUMBL) to assess pathway-level differences between these functional classes.

### Expression analysis and transcriptional dynamics in PDMR models

The expression levels of NUMB and NUMBL isoforms were analysed across different tumour passages available in the PDMR repository (Resection, P0–P4). To this end, the samples were previously annotated using a metadata file associated with the quantification files, which included information on the patient identifier, tumour passage, Oncotree code, and diagnostic classification. Isoform expression dynamics across tumour passages were visualized using normalized bar plots (relative to the Resection condition). To explore functional associations between NUMB/NUMBL isoforms and key pathways in cancer, Pearson correlation coefficients were calculated between each isoform and transcripts annotated in the Notch, Hippo, WNT, and SHH pathways. This analysis was restricted to the Resection, P0, and P4 passages. The results were represented using *pheatmap*, with a scale centred at zero (red colours for positive correlations, blue for negative ones).

To further characterize pathway-level transcriptional dynamics across tumour passages, differential expression was computed between consecutive stages (P0 vs Resection, P1 vs P0, P2 vs P1, P3 vs P2, and P4 vs P3) using *limma [95]*, followed by pathway enrichment and active subnetwork analysis with *pathfinder* (RRID:SCR_025459) [96]. For selected cancer-related pathways, genes identified across all comparisons were aggregated, mean log2FC were calculated per transition, and cumulative log2FC values were obtained by sequential summation starting from the Resection condition. This strategy enabled reconstruction of pathway-specific transcriptional trajectories across tumour passages and quantification of pathway reactivation between early and late PDX stages.

To characterize the global transcriptional organization during tumour progression, UMAP dimensionality reduction was applied to the gene expression profiles belonging to each of the four pathways analysed, using the *uwot* package (version 0.2.3, CRAN) [97]. The expression matrix was transposed to represent each sample as an observation, and UMAP was applied with a Euclidean metric, n_neighbors = 15, and min_dist = 0.1. Subsequently, an unsupervised clustering analysis using k-means was performed on the embedded UMAP space. The optimal number of clusters was determined using the elbow method and the average silhouette index.

Finally, alluvial plots were generated to visualize the transition of tumour samples between passages and clusters defined using UMAP + k-means, illustrating conserved or divergent transcriptional trajectories. To do this, the number of samples per passage and cluster combination was counted, and frequencies were plotted using *ggalluvial* (RRID:SCR_021253) [98]. The clusters obtained using k-means were manually renamed to improve visual interpretation, grouping those with similar dynamics together and facilitating the tracking of transitions between tumour passages in the alluvial plot.

### Functional and clinical analysis based on the *NUMB-score*

To investigate the functional impact of NUMB and NUMBL isoform profiles, a functional score (*NUMB-score*) was defined based on their expression levels in breast cell lines obtained from the CCLE. The score was calculated as the difference between the mean expression of potentially oncogenic isoforms (p72/p71) and the mean of suppressor isoforms (p66/p65/NUMBL). To reduce ambiguity near the centre of the distribution, the 10% of cell lines with values closest to the median score were excluded, dividing the remaining lines into two groups: high and low NUMB. A differential expression analysis was then performed between the two groups using the *limma* package (RRID: SCR_010943) [95], incorporating the score category as an explanatory variable. A model was built directly representing the expression means of each group, compared using a linear contrast defined as high − low. The results were FDR-adjusted using the Benjamini-Hochberg method, and genes with |log2FC| > 1 were selected. This set was used to perform functional enrichment analysis with the *enrichGO()* function of the *clusterProfiler* package, using Gene Ontology (Biological Process) annotations and an adjusted significance threshold (q < 0.05).

To specifically evaluate the effect of *NUMB-score* on key signalling pathways in cancer, differentially expressed genes belonging to the Notch (hsa04330), WNT (hsa04310), SHH (hsa04340), and Hippo (hsa04390) pathways were selected, based on annotations obtained from the KEGG database (RRID: SCR_012773). Those with |log2FC| > 0.8 and p-value < 0.05 were considered significant, and their expression levels were compared between the high and low *NUMB-score* groups.

The association between *NUMB-score* and drug sensitivity was also evaluated. Drug response data (AUC and log(IC50)) from breast cell lines available in the GDSC1 and GDSC2 studies [99, 100], downloaded from the Genomics of Drug Sensitivity in Cancer portal (RRID:SCR_011956, https://www.cancerrxgene.org), were integrated. For each drug, mean values were calculated in the high and low groups, and Student’s t tests were applied to evaluate differences in sensitivity. The results were visualized using volcano plots, highlighting the compounds with the greatest difference in efficacy between groups.

Independently, the *NUMB-score* was also calculated in the TCGA-BRCA set samples, using the pre-normalized expression levels of the same isoforms. This allowed examining their distribution in different clinical and molecular contexts. Comparisons included: (i) tumour status (tumour vs. non-tumour), (ii) AJCC clinical stage, and (iii) PAM50 molecular subtypes. Non-tumour samples were used as a reference. Statistical comparisons of *NUMB-score* distributions between groups (tumour status, clinical stage, or molecular subtype) were performed using the Wilcoxon test.

Finally, to assess the prognostic value of the *NUMB-score* in breast cancer, clinical data from the TCGA-BRCA project were integrated with the gene expression matrix and the previously calculated score, considering only tumour samples. Survival time was defined as the days elapsed from diagnosis to death or, in living patients, to the date of last follow-up. In cases where this information was unavailable, an artificial censor value of 4500 days was assigned to allow patient inclusion without introducing bias into the risk analysis. A single cutoff point was established for the *NUMB-score* (threshold = 2.55), classifying patients into two groups: High Score (≥ 2.55) and Low Score (< 2.55). The association between the score and overall survival was evaluated using Kaplan-Meier curves, compared using the log-rank (χ²) test. Survival analysis was conducted using the R packages *survival* (RRID: SCR_021137) and *survminer* (RRID: SCR_021094) [101, 102]. For the NUMB-score analysis, robustness was assessed by influence diagnostics based on DFBETA residuals from a Cox proportional hazards model, and survival analyses were repeated after exclusion of the top 1% most influential cases. In addition, isoform-specific Kaplan–Meier analyses were performed using expression-based stratification, with optimal cutoffs determined independently for each isoform by a scanning procedure enforcing a minimum group size of 15% of the cohort.

## Supporting information

Supplementary Table 2. Pathway-associated genes and transcript-level log2FC values

Supplementary Table 3. KEGG GSEA results for shared terms

Supplementary Table 4. Gene expression values underlying Figure 3F

Supplementary Table 5. Expression values underlying Figure 5D

Supplementary Table 6. Log-transformed IC50 values for Figure 5E

Supplementary Table 7. Sample-level traceability across TCGA-BRCA, CCLE, and PDMR

## DECLARATIONS

### ETHICS APPROVAL AND CONSENT TO PARTICIPATE

Not applicable.

### CONSENT FOR PUBLICATION

Not applicable.

### AVAILABILITY OF DATA AND MATERIAL

All datasets analysed in this study are publicly available. TCGA-BRCA RNA-seq isoform expression and clinical data were obtained from the NCI Genomic Data Commons (GDC; RNA-seq Level 3 RSEM isoform-normalized data dated 2015-08-06). CCLE expression data were obtained from the Cancer Cell Line Encyclopedia (CCLE; RNA-seq transcript TPM file dated 2018-09-29), and drug-response data were integrated with GDSC (release dated 2023-10-27) after harmonising cell-line identifiers. Breast PDX isoform quantifications and metadata were obtained from the PDMR resource. Key analysis outputs and sample-level annotations, including transcript-level differential expression, pathway enrichment results, metabolic expression matrices, drug-response statistics, and *NUMB-score* assignments, are provided in the Supplementary Information files (Supplementary Tables 2–7).

### CODE AVAILABILITY

All custom scripts used for data processing and analysis are available in a GitHub repository. A permanent archived version of the code will be deposited in Zenodo and the corresponding DOI will be provided upon acceptance.

### COMPETING INTERESTS

The authors declare that they have no competing interests.

## FUNDING

This research was supported by grants from Ministerio de Ciencia, Innovación y Universidades (MCIU) Plan Estatal de I+D+I 2018, Agencia Estatal de Investigación (AEI) and Fondo Europeo de Desarrollo Regional (MCIU/AEI/FEDER, UE): PID2024-155394OB-I00 and PID2021-122629OB-I00 funded by MCIN/AEI/10.13039/501100011033 and by “ERDF A way of making Europe”, by the “European Union”; AEI-MICIU/FEDER (iDIFFER network: RED2024-153635-T). Additional grants from CIBER de Cáncer (CB16/12/00275), from Consejeria de Salud (PI-0397-2017) and Projects P18-RT-2501; and DGP_PIDI_2024_00907 from the Regional Ministry of Economic Transformation, Industry, Knowledge and Universities. Junta de Andalucía. Special thanks to the AECC (Spanish Association of Cancer Research) Founding Ref. GC16173720CARR for supporting this work.

## ACKNOWLEDGEMENTS

Special thanks to the AECC Foundation for supporting this work.

## AUTHORS CONTRIBUTION

AC and JMG-H conceived and designed this study. JMG-H and SO-C performed bioinformatic analysis. AC and JMG-H drafted, edited and revised the manuscript. All authors approved the manuscript.

## SUPPLEMENTARY MATERIAL

### Supplementary data

**Supplementary Table S1.** Abbreviations used in Figure 1.

| Category | Abbreviation | Full name |
| --- | --- | --- |
| TCGA tumour type | ACC | Adrenocortical carcinoma |
|  | BLCA | Bladder urothelial carcinoma |
|  | BRCA | Breast invasive carcinoma |
|  | CESC | Cervical squamous cell carcinoma and endocervical adenocarcinoma |
|  | CHOL | Cholangiocarcinoma |
|  | COAD | Colon adenocarcinoma |
|  | DLBC | Diffuse large B-cell lymphoma |
|  | ESCA | Esophageal carcinoma |
|  | GBM | Glioblastoma multiforme |
|  | HNSC | Head and neck squamous cell carcinoma |
|  | KICH | Kidney chromophobe |
|  | KIRC | Kidney renal clear cell carcinoma |
|  | KIRP | Kidney renal papillary cell carcinoma |
|  | LAML | Acute myeloid leukemia |
|  | LGG | Lower grade glioma |
|  | LIHC | Liver hepatocellular carcinoma |
|  | LUAD | Lung adenocarcinoma |
|  | LUSC | Lung squamous cell carcinoma |
|  | OV | Ovarian serous cystadenocarcinoma |
|  | PAAD | Pancreatic adenocarcinoma |
|  | PCPG | Pheochromocytoma and paraganglioma |
|  | PRAD | Prostate adenocarcinoma |
|  | READ | Rectum adenocarcinoma |
|  | SARC | Sarcoma |
|  | SKCM | Skin cutaneous melanoma |
|  | STAD | Stomach adenocarcinoma |
|  | TGCT | Testicular germ cell tumors |
|  | THCA | Thyroid carcinoma |
|  | THYM | Thymoma |
|  | UCEC | Uterine corpus endometrial carcinoma |
|  | UCS | Uterine carcinosarcoma |
| AS event type | AA | Acceptor site |
|  | AD | Alternative donor site |
|  | AP | Alternative promoter |
|  | AT | Alternative termination site |
|  | ES | Exon skipping |
|  | IR | Intron retention |
|  | MES | Mutually exclusive exons |

**Supplementary Table 2.** Pathway-associated genes and transcript-level log2FC values used in the isoform-specific analyses. Each row corresponds to an individual transcript. Consequently, genes with multiple annotated transcripts may appear more than once, reflecting transcript- and isoform-specific regulatory differences rather than aggregated gene-level effects.

**Supplementary Table 3.** KEGG GSEA results for shared terms across NUMB/NUMBL isoform groups in TCGA-BRCA.

**Supplementary Table 4.** Gene expression values underlying the metabolic heatmap shown in Figure 3F.

**Supplementary Table 5**. Expression values and transcript identifiers underlying Figure 5D.

**Supplementary Table 6.** Log-transformed IC50 values and statistical association metrics for all compounds tested in CCLE breast cancer cell lines, comparing high and low NUMB-score groups, corresponding to the drug-response analysis shown in Figure 5E.

**Supplementary Table 7.** Sample-level traceability across TCGA-BRCA, CCLE breast cancer cell lines, and PDMR (PDX) models.

**Supplementary Figure 1.**
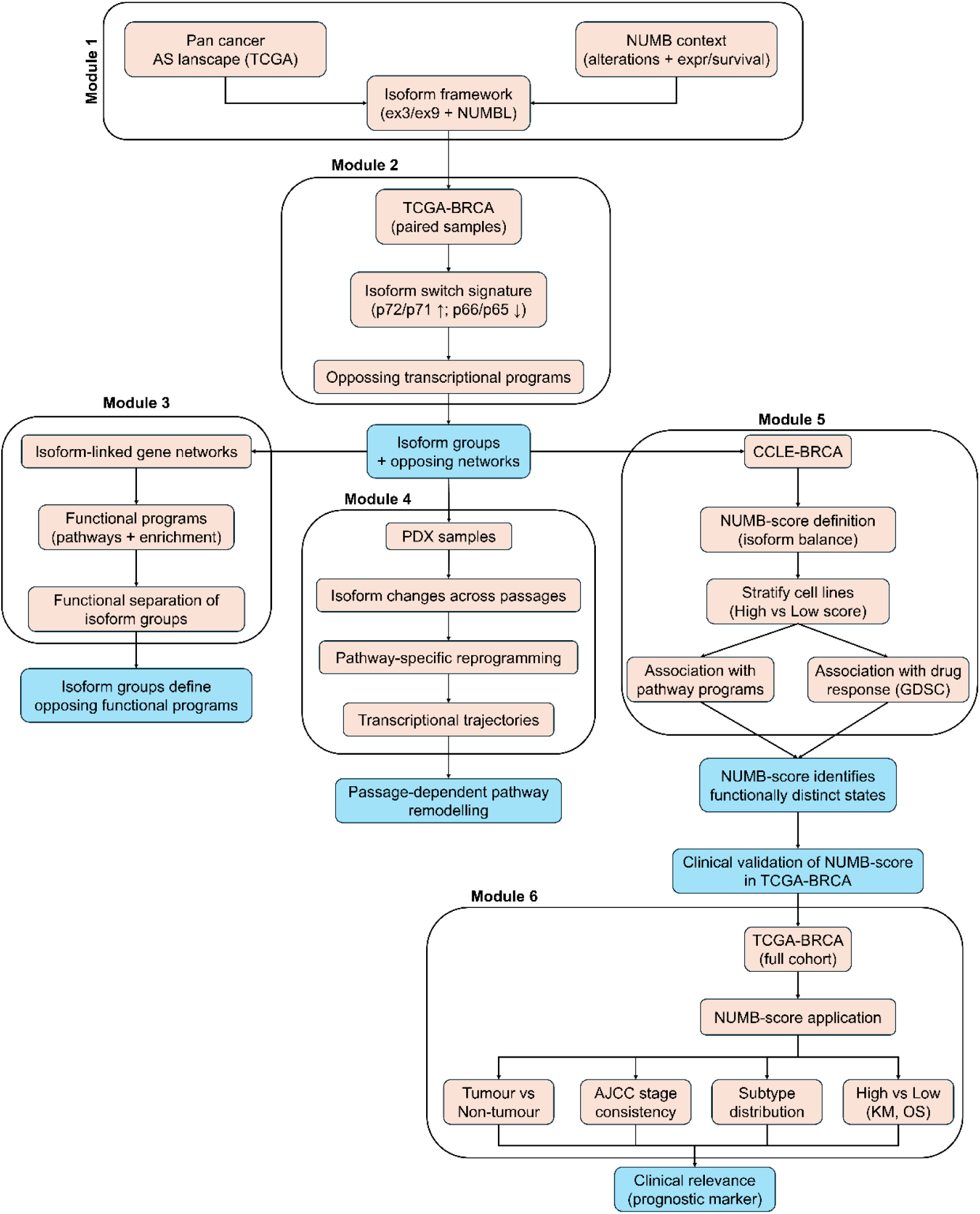
Overview of the study workflow and computational analyses. The diagram summarizes the sequential datasets and analysis steps used throughout the study, from pan-cancer TCGA alternative splicing profiling and the definition of a NUMB isoform framework (exon 3/exon 9 inclusion and NUMBL), to TCGA-BRCA paired tumour/non-tumour analyses defining the NUMB isoform switch signature and associated transcriptional programs. NUMB isoform groups are linked to gene networks and functional pathways, and isoform dynamics are evaluated across passages in patient-derived xenograft (PDX) samples. A NUMB-score based on isoform balance is then defined in CCLE-BRCA, used to stratify cell lines into functionally distinct states, and associated with pathway programs and drug response patterns (GDSC). Finally, the NUMB-score is clinically validated in the full TCGA-BRCA cohort, including associations with tumour status, AJCC stage, molecular subtypes and overall survival (KM/OS), supporting its clinical relevance as a prognostic marker.

**Supplementary Figure 2.**
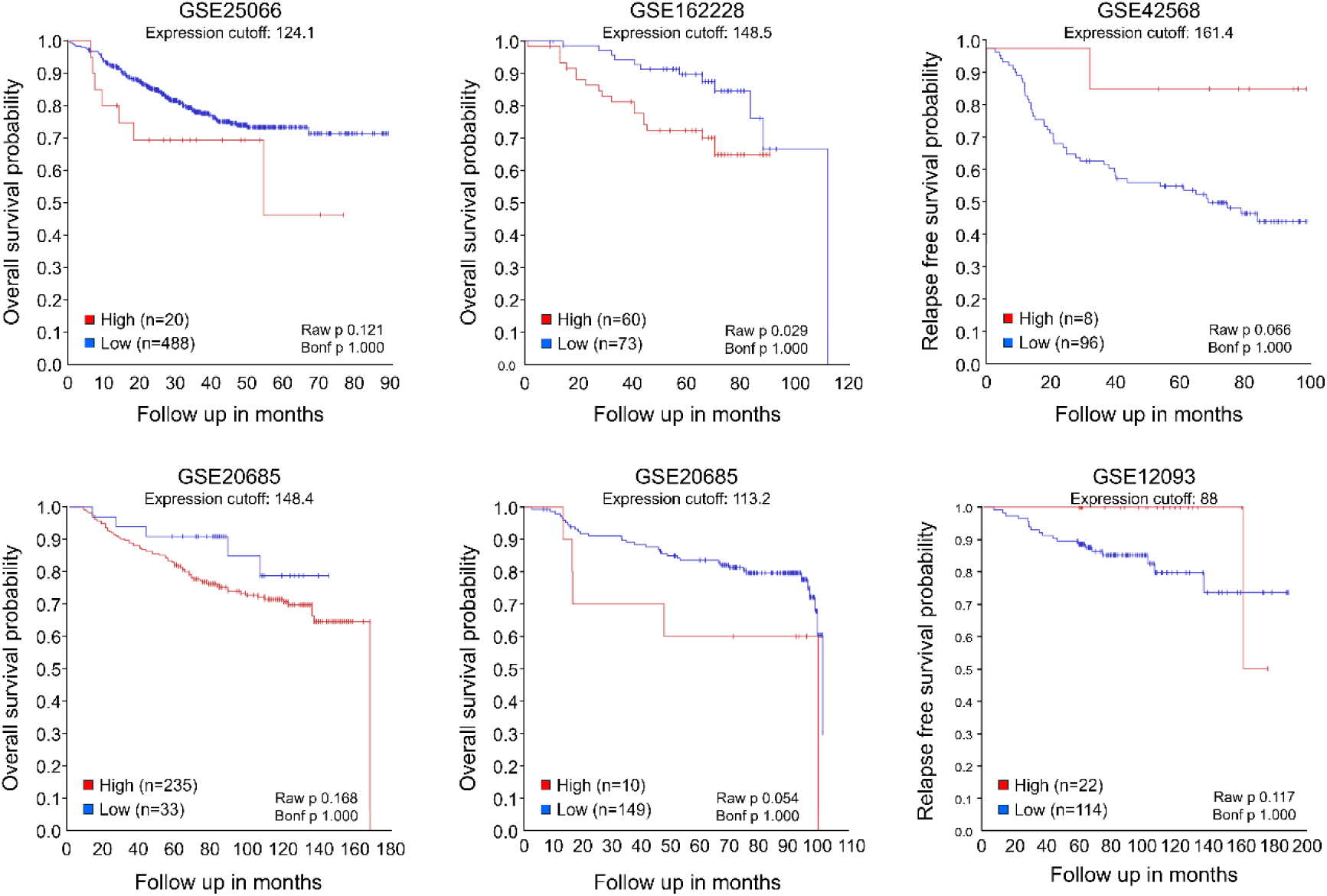
*NUMB* expression behaves as a marker of poorer or better survival in breast cancer. Kaplan–Meier plots from independent breast cancer cohorts show that lower *NUMB* expression is associated with better survival in GSE25066, GSE162228, GSE20685 and GSE1456, whereas it is associated with worse survival in GSE42568 and GSE12093.

**Supplementary Figure 3.**
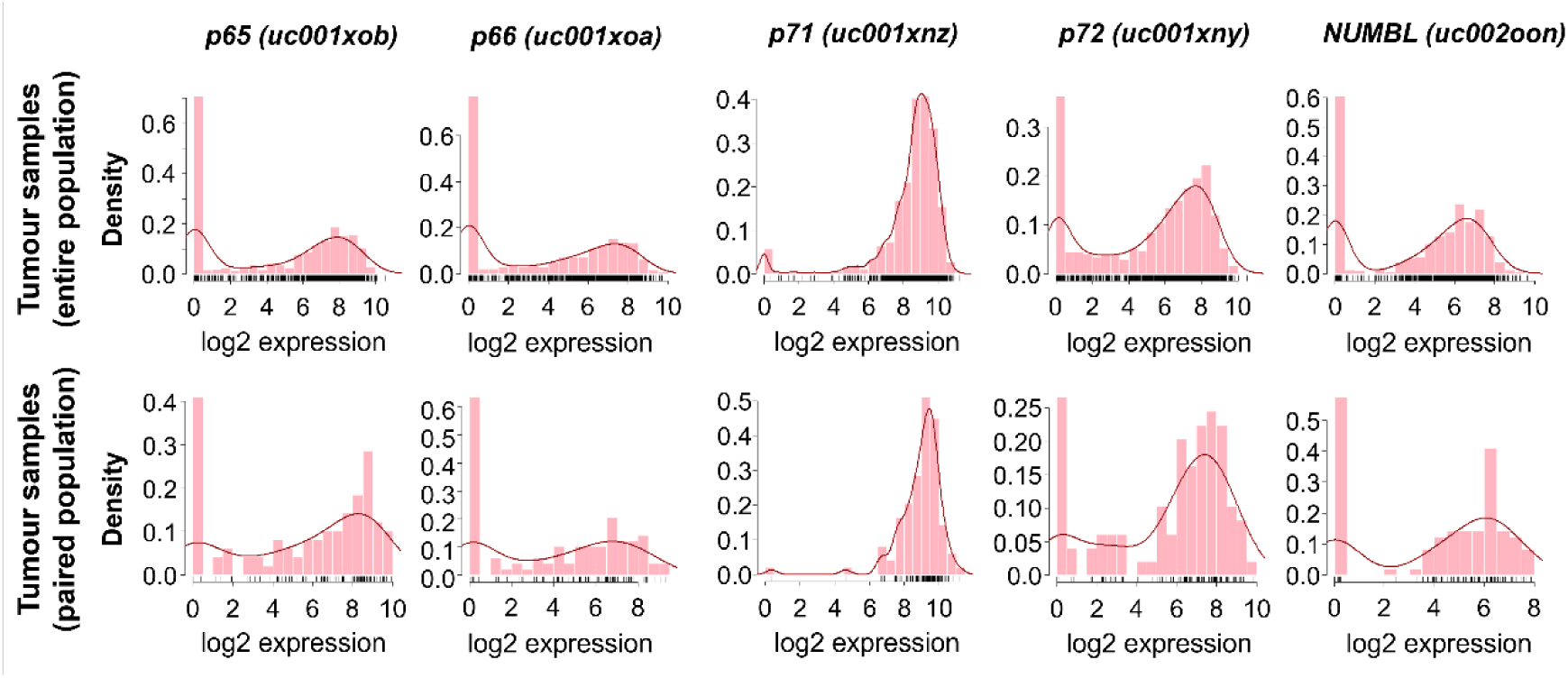
Histograms showing the expression distribution of the *NUMB* isoforms and *NUMBL* in tumour samples. The upper row shows data from the entire population, while the lower row corresponds to tumour samples paired to non-tumour. The similarity in distribution suggests that the paired subset can be considered a representative group of the entire population.

**Supplementary Figure 4.**
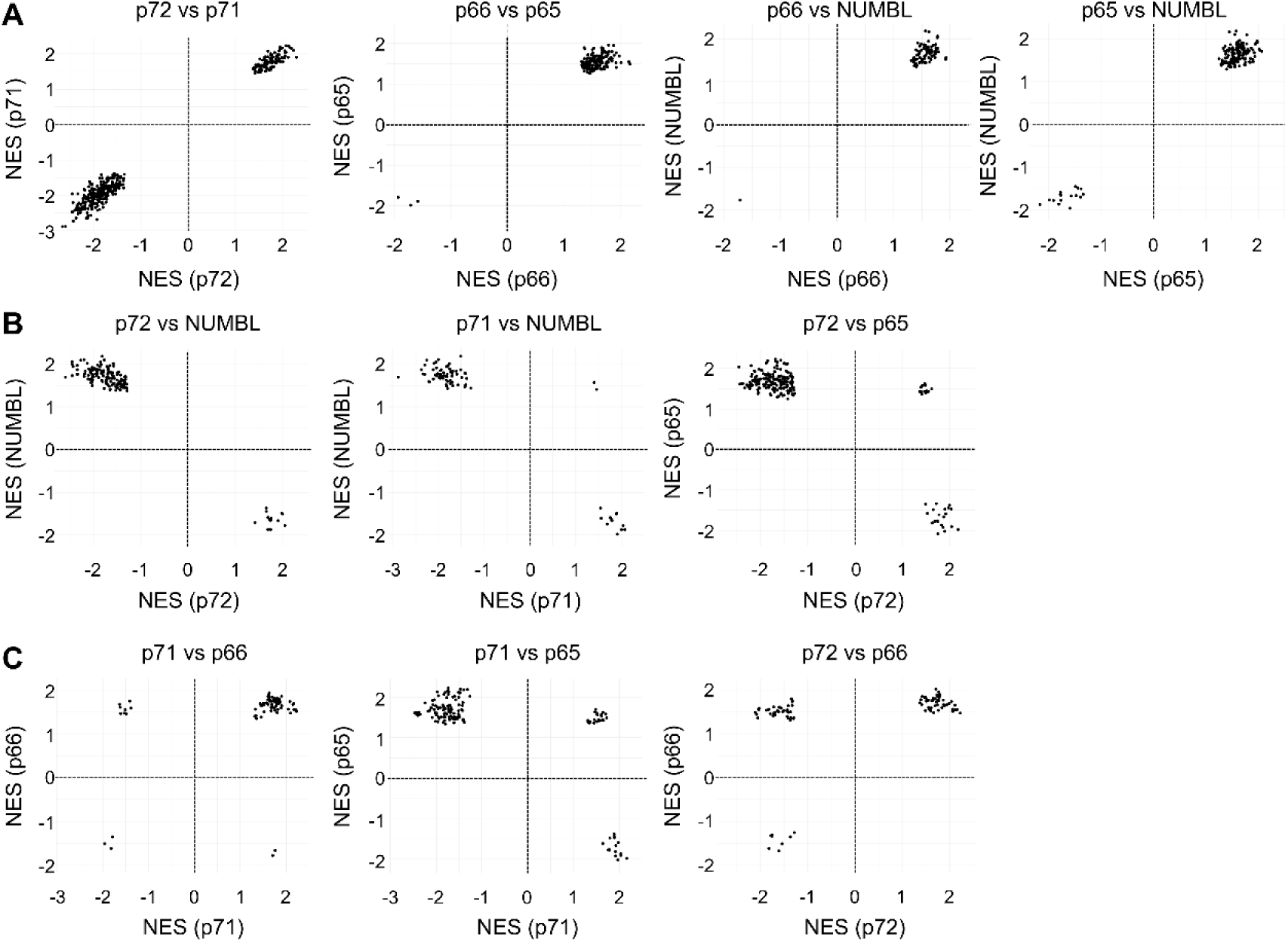
Scatter plots comparing Normalized Enrichment Scores (NES) obtained from GSEA analyses between NUMB isoforms and NUMBL. **A)** The isoform pairs p72/p71, p66/p65, p66/NUMBL, and p65/NUMBL displayed highly similar NES values across all shared GO terms analysed by GSEA. **B)** The isoform pair p72/NUMBL showed completely opposite NES values, suggesting distinct gene set associations for each isoform. A similar pattern was observed for p71/NUMBL and p72/p65, although some GO terms still showed concordant positive NES values. **C)** The isoform pairs p71/p66, p71/p65, and p72/p66 exhibited a more mixed pattern, suggesting partial convergence in pathway associations among these isoforms.

**Supplementary Figure 5.**
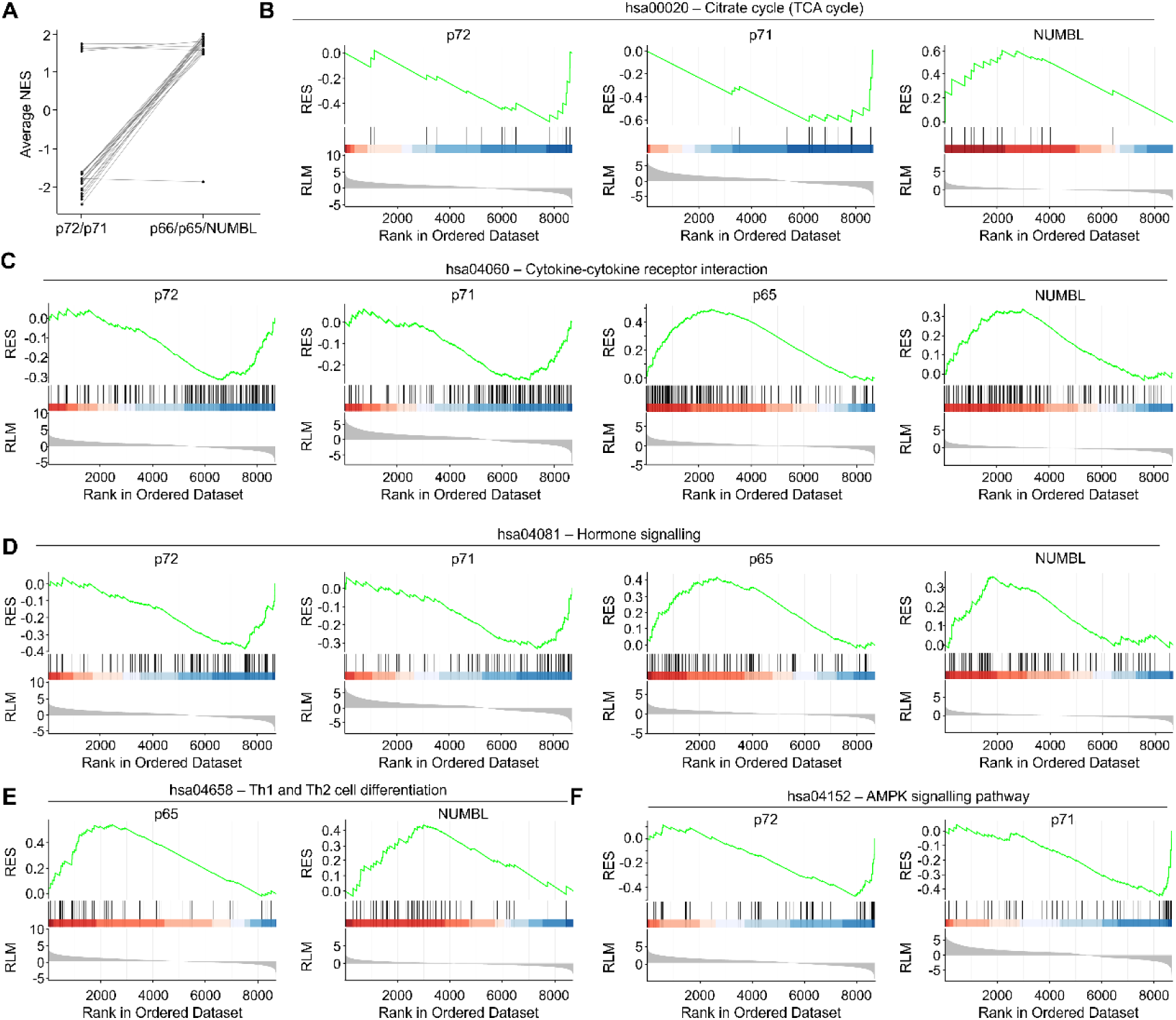
GSEA analysis shows functional divergence between NUMB isoforms in breast cancer. **A)** Comparison of NES values for shared KEGG terms across NUMB isoform groups. Each line connects the average NES value for the same KEGG term between the p72/p71 and p66/p65/NUMBL isoforms. A systematic reversal of the enrichment direction (from negative to positive) is observed, indicating opposing regulation of key biological pathways depending on the expressed isoform. **B)** The opposing association of the citrate cycle (hsa00020) with NUMB isoforms, shown by a negative enrichment trend for p72 and p71 and a positive association with NUMBL, suggests a lower dependence on mitochondrial metabolism when p72/p71 isoforms are predominantly expressed. **C)** The cytokine-cytokine receptor interaction pathway (hsa04060) shows negative enrichment in p72/p71-associated profiles and positive enrichment in p65/NUMBL-associated profiles, indicating an opposite association with this immune pathway depending on the predominant isoform. **D)** The hormonal signalling pathway (hsa04081) is negatively associated with p72/p71 and positively with p65/NUMBL, reflecting an inverse correlation with the expression of these isoforms in relation to hormonal processes. **E)** The Th1 and Th2 differentiation pathway (hsa04658) is exclusively enriched in profiles correlated with p65 and NUMBL, suggesting a specific link with active immune functions. **F)** The AMPK signalling pathway (hsa04152), typically associated with tumour suppressor functions, shows negative enrichment only in association with the p72 and p71 isoforms. RES: Running Enrichment Score; RLM: Rank List Metric.

**Supplementary Figure 6.**
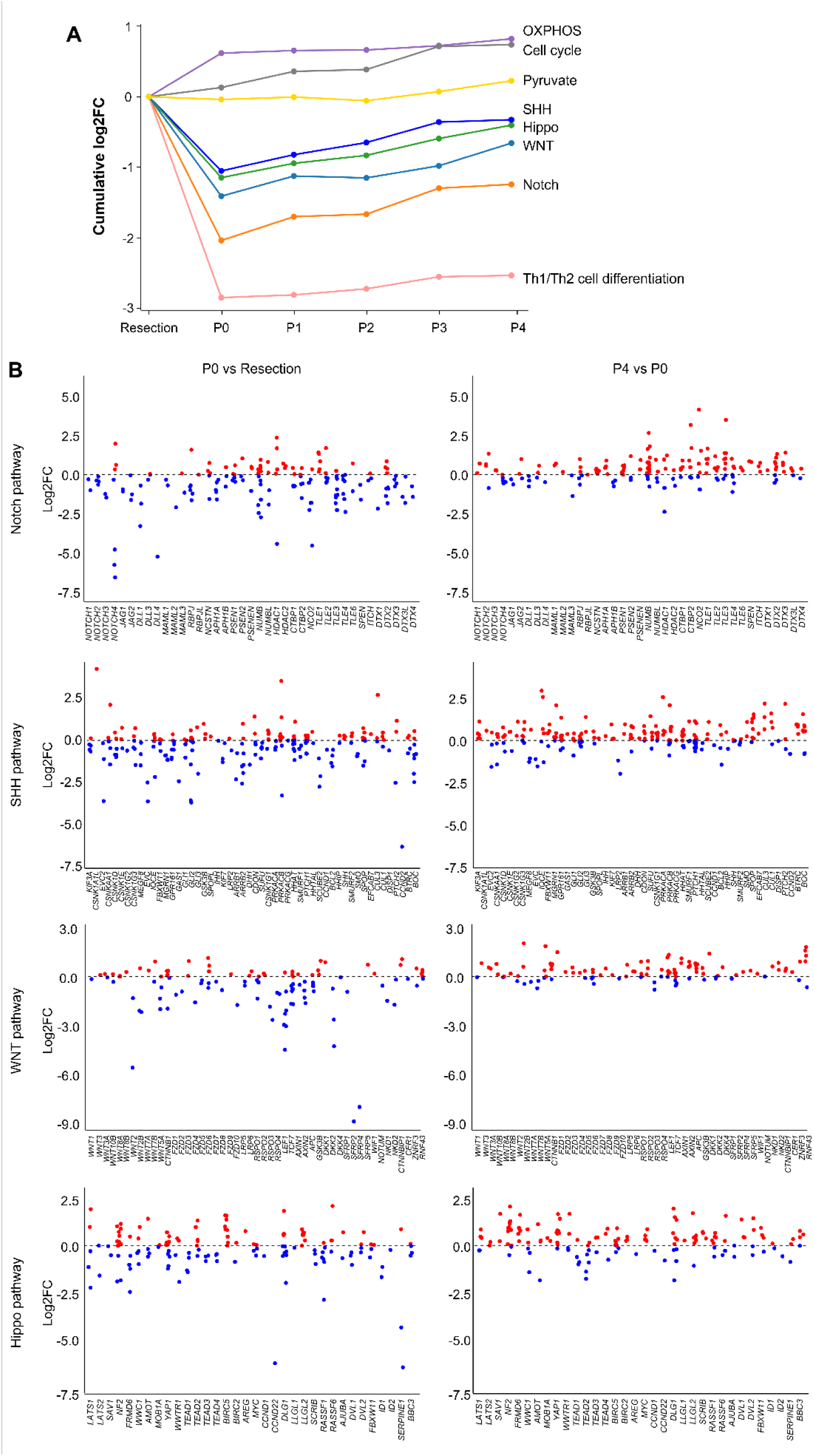
Dynamic transcriptional remodelling of key signalling pathways in PDX tumours. **A)** Cumulative log2FC in gene expression for multiple biological pathways across serial PDX passages compared to Resection. After implantation in mice, most signalling pathways show a marked downregulation, whereas pathways such as OXPHOS and cell cycle genes increase progressively with each passage, indicating a shift in tumour transcriptional programs. **B)** Gene-level expression changes for individual isoforms in representative signalling pathways (Notch, SHH, WNT, and Hippo). For each pathway, isoform expression is compared between P0 vs Resection (left plots) and P4 vs P0 (right plots). After implantation (P0), a general suppression of key genes is observed (blue dots), indicating early transcriptional reprogramming upon xenografting. Over subsequent passages (P4), partial recovery of gene expression is evident (red dots), suggesting progressive reactivation of these signalling pathways in the evolving PDX environment.

**Supplementary Figure 7.**
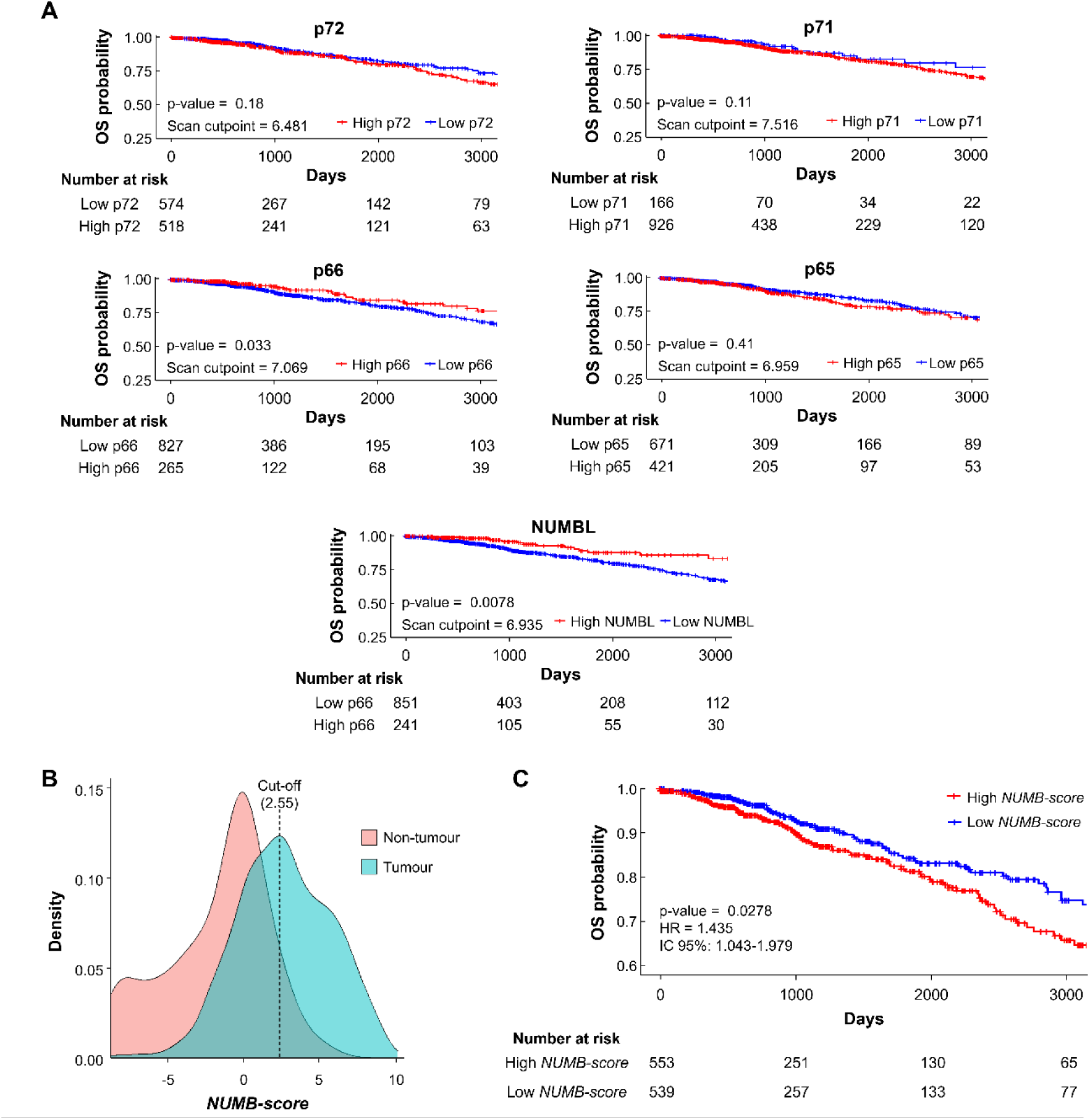
Isoform-level survival analyses and NUMB-score distribution in the TCGA-BRCA cohort. **A)** Kaplan–Meier overall survival curves for individual NUMB isoforms (p72, p71, p66, p65) and NUMBL in the TCGA-BRCA cohort, using scan-derived cutpoints with a minimum group size constraint. p66 and NUMBL show significant associations with survival, whereas p72, p71, and p65 do not. Numbers at risk are shown below each plot. **B)** Density distribution of *NUMB-score* values in tumour and non-tumour samples from TCGA-BRCA. The dashed line indicates the cut-off used to stratify samples. **C)** Kaplan–Meier overall survival analysis based on NUMB-score stratification in the full TCGA-BRCA cohort (without removal of influential samples), showing reduced survival in the high *NUMB-score* group. Hazard ratio (HR), 95% confidence interval (CI), and log-rank p-value are indicated.

## Notes

### Competing Interest Statement

The authors have declared no competing interest.

